# A functional screen for ubiquitin regulation identifies an E3 ligase secreted by *Pseudomonas aeruginosa*

**DOI:** 10.1101/2024.09.18.613774

**Authors:** Cameron G. Roberts, Supender Kaur, Aaron J. Ogden, Michael E. Divine, Gus D. Warren, Donghoon Kang, Natalia V. Kirienko, Paul P. Geurink, Monique P.C. Mulder, Ernesto S. Nakayasu, Jason E. McDermott, Joshua N. Adkins, Alejandro Aballay, Jonathan N. Pruneda

## Abstract

Ubiquitin signaling controls many aspects of eukaryotic biology, including targeted protein degradation and immune defense. Remarkably, invading bacterial pathogens have adapted secreted effector proteins that hijack host ubiquitination to gain control over host responses. These ubiquitin-targeted effectors can exhibit, for example, E3 ligase or deubiquitinase activities, often without any sequence or structural homology to eukaryotic ubiquitin regulators. Such convergence in function poses a challenge to the discovery of additional bacterial virulence factors that target ubiquitin. To overcome this, we have developed a workflow to harvest natively secreted bacterial effectors and functionally screen them for ubiquitin regulatory activities. After benchmarking this approach on diverse ligase and deubiquitinase activities from *Salmonella* Typhimurium, Enteropathogenic *Escherichia coli*, and *Shigella flexneri*, we applied it to the identification of a cryptic E3 ligase activity secreted by *Pseudomonas aeruginosa*. We identified an unreported *P. aeruginosa* E3 ligase, which we have termed *Pseudomonas* Ub ligase 1 (PUL-1), that resembles none of the other E3 ligases previously established in or outside of the eukaryotic system. Importantly, in an animal model of *P. aeruginosa* infection, PUL-1 ligase activity plays an important role in regulating virulence. Thus, our workflow for the functional identification of ubiquitin-targeted effector proteins carries promise for expanding our appreciation of how host ubiquitin regulation contributes to bacterial pathogenesis.

## INTRODUCTION

Signaling networks mediated by the post-translational modifier ubiquitin regulate a vast domain of eukaryotic biology. Through a process termed ubiquitination, the 76-amino acid protein ubiquitin (Ub) is conjugated via its C-terminus onto target proteins, typically at lysine (Lys, K) residues, to trigger signaling outcomes such as trafficking or degradation. The diversity in Ub-dependent signaling outcomes arises, in part, from the extension of polymeric Ub (polyUb) chains that act as distinct signaling molecules. For example, polyUb chains linked via K48 form the classical signal for proteasomal degradation, while those linked via K63 are non-degradative and facilitate protein recruitment in pathways such as endocytosis, immune activation, or the DNA damage response^1,2^. Ub conjugation is heavily regulated by a cascade of E1 Ub-activating, E2 Ub-conjugating, and E3 Ub ligase enzymes, while deconjugation is mediated by deubiquitinases (DUBs). E3 ligases represent the largest set of Ub regulators, with over 600 examples in humans. With few exceptions^3,4^, human E3 ligases fall into roughly three structurally and mechanistically distinct families: Really Interesting New Gene (RING) ligases^5^, Homologous to E6AP C-terminus (HECT) ligases^6^, and RING-between-RING (RBR) ligases^7^. While RING ligases catalyze direct Ub conjugation from an E2∼Ub intermediate onto a substrate, HECT and RBR ligases form a final E3∼Ub intermediate, in which the Ub C-terminus is activated onto an active site cysteine (Cys, C) through a high-energy thioester linkage. These three mechanisms of Ub conjugation are highly conserved across eukaryotic biology.

Remarkably, despite lacking a canonical Ub system of their own, pathogenic viruses and bacteria have evolved virulence factors capable of regulating Ub signaling of their eukaryotic hosts^8,9^. Bacteria, in particular, have evolved a multitude of strategies to redirect, block, or eliminate host Ub signaling through the action of secreted ‘effector’ proteins^10^. While some of these strategies of Ub regulation mimic those used by eukaryotes, others appear to be entirely distinct and likely arose through convergent evolution^8^. Bacterial E3 ligases, for example, fall into at least seven distinct families, only one of which resembles eukaryotic RING ligases. Stemming from this convergent nature, the identification of bacterial Ub regulators has been cumbersome and limited to select pathogens. Previous efforts have relied upon labor-intensive approaches, such as individual mapping of protein-protein interactions^11,12^, or introduce a heavy bias toward known enzymes and mechanisms, such as sequence/structural homology or the use of activity-based probes (ABPs)^13–15^. On the other hand, a functional approach focused on bacterial DUBs specific to linear (M1-linked) polyUb chains identified an effector protein from *Legionella pneumophila* that is distinct from any DUB families previously observed in eukaryotes, viruses, or bacteria^16^. An unbiased discovery approach, therefore, has the potential to identify unprecedented Ub regulators that mediate bacterial virulence.

From the animal and plant pathogens that have been studied thus far, it has become increasingly clear that subversion of host Ub signaling is a commonly adopted virulence strategy. The gastrointestinal pathogen *Shigella flexneri*, for example, has devoted nearly one-third of its effector repertoire toward regulating the Ub system^17^. Still, other medically relevant pathogens, with elaborate mechanisms of host manipulation, lack any described methods to regulate host ubiquitination. Among them, *Pseudomonas aeruginosa* represents a critical target for developing an improved understanding of host-pathogen interactions. *P. aeruginosa* is an opportunistic, Gram-negative pathogen that poses a severe health risk for immunocompromised individuals and is among the leading causes of nosocomial infections. It is also of extreme concern for antimicrobial resistance, as highly resistant strains of *P. aeruginosa* were associated with over 300,000 deaths worldwide in the year 2019 alone, representing a massive and increasing health burden^18^. *P. aeruginosa* encodes a large repertoire of virulence factors, including a Type III Secretion System (T3SS) known to secrete a combination of ExoU, ExoS, ExoT, and ExoY effector proteins into the host cell^19^. While ExoT levels are regulated by host ubiquitination and ExoU phospholipase activity requires Ub as a cofactor^20,21^, no *P. aeruginosa* effectors have been identified to directly regulate host Ub signaling. Furthermore, no Ub regulators are readily apparent from sequence and structural homology analyses, raising question as to whether *P. aeruginosa* has not adapted strategies to regulate ubiquitination, or if cryptic Ub regulators remain to be discovered.

Herein, we developed a novel unbiased method to survey Ub regulatory activities of natively secreted bacterial effectors. By stimulating secretion in culture, pools of bacterial effector proteins could be harvested as input for highly sensitive, fluorescence-based assays of Ub regulatory activities. In this manner, secreted ligase and DUB activities were observable from *Salmonella* Typhimurium, enteropathogenic *Escherichia coli* (EPEC), and *S. flexneri*, consistent with previous reports^22–25^. Applying this approach to *P. aeruginosa*, we identified a cryptic E3 ligase activity from the gene product PA2552, which we have renamed to *Pseudomonas* Ub ligase 1 (PUL-1). *In vitro*, PUL-1 catalyzes a mixture of mono- and polyubiquitination downstream of human E1 and E2 enzymes. Interestingly, PUL-1 represents an uncharacterized family of Cys-based E3 ligases that are conserved among *P. aeruginosa* clinical isolates as well as other bacterial pathogens. *In vivo*, the ligase activity of PUL-1 modulates *P. aeruginosa* virulence in a *C. elegans* model of infection. Thus, the approach we describe offers a straightforward and unbiased opportunity to identify cryptic Ub regulators secreted as virulence factors by bacterial pathogens.

## RESULTS

### A functional screen for ubiquitin regulation

To overcome the challenge of identifying evolutionarily convergent bacterial effectors that regulate ubiquitination, we sought to implement an unbiased functional approach. As sensitive and robust measures for Ub conjugation and deconjugation are readily available, the primary obstacle was isolation of candidate bacterial effector molecules. To eliminate contributions from the eukaryotic Ub regulatory system that would come with bacterial infections, we instead utilized established conditions that artificially stimulate effector secretion in bacterial culture. In many cases, these stimulatory conditions mimic native cues sensed by the bacterial secretion systems, such as lowered pH that triggers the *S.* Typhimurium SPI-II T3SS in the context of an acidified vacuole^26^. Following stimulation, secreted effectors could be harvested from the culture supernatant, concentrated by native ammonium sulfate precipitation, and resolubilized for biochemical activity assays (**Fig. 1A**). Meanwhile, a bacterial lysate could be prepared from the culture pellet, for comparison to the secreted protein fraction (**Fig. 1A**). As an alternative to stimulation, strains with mutations in secretion regulators, leading to constitutive secretion, can be used in a similar workflow. *S. flexneri*, EPEC, and *S.* Typhimurium were selected for proof-of-principle experiments, as *S.* Typhimurium encodes a reported DUB^22^, and all three encode reported E3 ligases^23–25^. *S. flexneri* effector secretion can be triggered with Congo Red or with the constitutively-secreting *ΔipaD* mutant^27,28^, EPEC secretion can be triggered by pH and salt conditions commonly present in Dulbecco’s modified Eagle’s medium (DMEM)^29^, while the SPI-I and SPI-II secretion systems of *S.* Typhimurium can be triggered with changes in salt or pH, respectively^26,30,31^ (**Fig. 1B**). Under these conditions, we could harvest natively secreted effector repertoires that were distinct from the lysate fractions, and unique to each bacterial secretion system (**Fig. 1B**).

**Figure 1:**
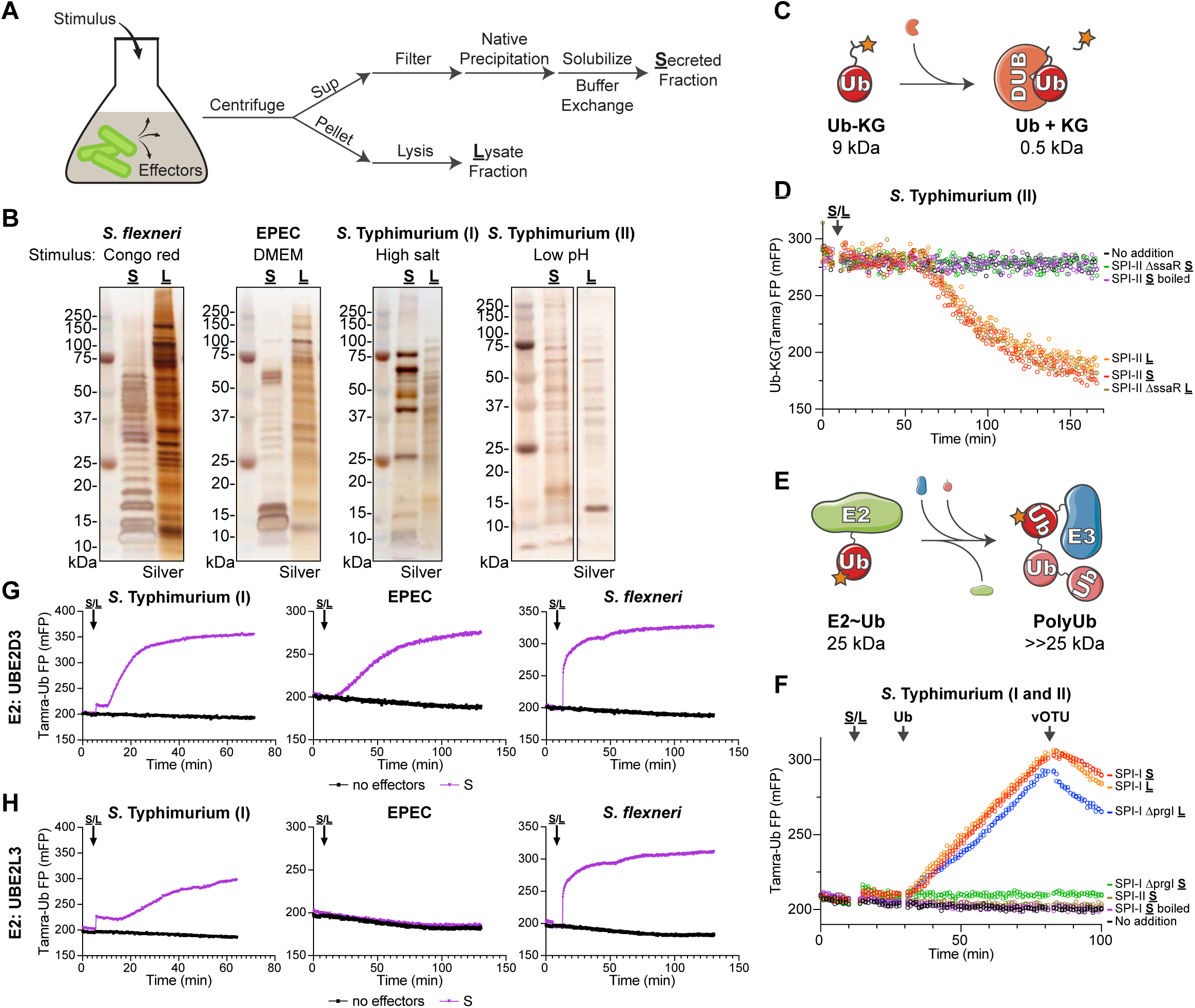
A functional screen for ubiquitin regulation. A. Schematic for the strategy used to stimulate effector secretion in bacterial culture prior to harvesting the secreted and lysate fractions. B. Silver-stained SDS-PAGE analysis of secreted (**<u>S</u>**) and lysate (**<u>L</u>**) fractions prepared following stimulation as indicated. C. Schematic for the FP-based assay for detection of DUB activity. D. Representative FP traces monitoring the Ub-KG(Tamra) DUB substrate following addition of the indicated pools of *S.* Typhimurium protein. E. Schematic for the FP-based assay for detection of E3 ligase activity. F. Representative FP traces monitoring the Tamra-Ub ligase substrate following addition of the indicated pools of *S.* Typhimurium protein. At the indicated timepoints, additional unlabeled Ub or the DUB vOTU were added to stimulate product extension or deconjugation, respectively. G. Representative FP traces monitoring the Tamra-Ub ligase substrate following addition of the indicated bacterial secreted fractions to reactions containing the promiscuous E2 enzyme, UBE2D3. H. As in **G)**, for reactions containing the Cys-specific E2 enzyme, UBE2L3.

Among the methods to detect Ub conjugation and deconjugation *in vitro*, those that utilize fluorescence polarization (FP) offer both broad applicability as well as high sensitivity. Deconjugation activity can be readily measured as a decrease in FP following cleavage of a Ub-KG(Tamra) substrate, in which Ub is natively isopeptide-linked to the ε-amino group of a fluorescent KG dipeptide^32^ (**Fig. 1C**). This substrate has been heavily used to characterize eukaryotic as well as bacterial DUBs^13,32,33^. The *S.* Typhimurium SPI-II secretion system reportedly delivers the DUB SseL into host cells^22^. Consistently, we observe cleavage of the Ub-KG(Tamra) substrate upon addition of the *S.* Typhimurium SPI-II secreted fraction (**Fig. 1D**). This activity is abolished upon boiling the secreted fraction, and is restricted to only the lysate fraction when prepared from a SPI-II secretion-deficient Δ*ssaR* mutant strain^34,35^ (**Fig. 1D**). Thus, our approach can readily detect natively secreted DUB activity among bacterial effectors.

Using a recently developed method called UbiReal^36,37^, Ub conjugation can also be monitored by FP. In this case, a fluorescently labeled Ub is conjugated through the E1-E2-E3 cascade, resulting in relative increases in molecular weight that coincide with increased FP (**Fig. 1E**). This approach has been effective in characterizing both eukaryotic and bacterial E3 ligases^36–38^. *S.* Typhimurium encodes four reported E3 ligases: the HECT-like effector SopA secreted by SPI-I and three NEL (novel E3 ligase) effectors secreted by SPI-II^23,25,39,40^. In a UbiReal assay that combines the two most promiscuous E2 enzymes^41^, UBE2D3 and UBE2L3, ligase activity consistent with SopA is observed in the SPI-I secreted fraction (**Fig. 1F**). Consistent with their reported auto-inhibited state in the absence of substrate^42^, we observe no ligase activity from NEL effectors in the SPI-II secreted fraction. The SPI-I secreted ligase activity can be reversed upon addition of the nonspecific DUB, vOTU^43^ (**Fig. 1F**). The activity is also ablated upon boiling of the secreted fraction, and is restricted to the SPI-I lysate fraction when prepared from a SPI-I secretion-deficient Δ*prgI* mutant strain^44,45^ (**Fig. 1F**). The E2348/69 strain of EPEC reportedly encodes an E3 ligase from the NleG family^24^, while *S. flexneri* reportedly encodes multiple NEL-family ligases^25^. Consistently, E3 ligase activity is observed in the EPEC and *S. flexneri* secreted fractions in UbiReal assays utilizing the highly promiscuous E2 UBE2D3 (**Fig. 1G**). However, while *S.* Typhimurium SopA and the *S. flexneri* NEL effectors utilize a Cys-dependent ligase mechanism, the EPEC NleG-type ligase utilizes a cysteine-independent mechanism akin to eukaryotic RING and U-box ligases. Accordingly, in UbiReal assays that incorporate the Cys-specific E2 UBE2L3, activity is only observed for *S.* Typhimurium SPI-I and *S. flexneri* secreted fractions (**Fig. 1H**). The E2- and cysteine-dependent nature of the secreted ligase activities can also be confirmed by conventional western blotting approaches (**Fig. S1A-C**). Thus, not only can natively secreted E3 ligase activities be observed with this approach, but the type of activity can be characterized as well.

### Detection of E3 ligase activity secreted by *P. aeruginosa*

With proof-of-principle for the functional approach established, we sought to screen for unidentified Ub regulatory activities. The opportunistic pathogen *P. aeruginosa* exploits a gamut of virulence mechanisms, including a T3SS, to infect a wide range of eukaryotic hosts. Despite several connections between its T3SS effectors and the host Ub system^20,21^, a mechanism of direct Ub regulation has not been identified. Using the PAO1 and PA14 reference strains of *P. aeruginosa*, we triggered effector secretion with an established calcium depletion approach^46^, and harvested the secreted and lysate fractions (**Fig. 2A**). These *P. aeruginosa* secreted fractions were notably more complex than those obtained from, e.g., *S. flexneri*, but were still distinct from the lysate fractions. Remarkably, introducing the PA14 secreted fraction into a UbiReal assay immediately revealed the presence of E3 ligase activity that could be reversed by addition of purified DUB (**Fig. 2B**). This ligase activity was also present in the PAO1 secreted fraction, and appeared to utilize a Cys-dependent mechanism, as activity was observed with both UBE2D3 and UBE2L3 (**Fig. 2C**). Further supporting a cysteine-based mechanism, the ligase activity was ablated following pre-treatment with either the Cys-reactive N-ethylmaleimide (NEM) or with Proteinase K (**Fig. 2D**). Consistent with other secreted virulence factors, the presence of ligase activity in the secreted fraction required calcium depletion (**Fig. 2D**). While the ligase activity was independent of the two-component system regulator GacA, its secretion did require the transcription regulator ExsA, which controls the T3SS regulon^47^ (**Fig. 2E, S2A**). Thus far, the repertoire of established T3SS effectors in PA14 is limited to ExoT, ExoU, and ExoY, yet the secreted fraction harvested from a Δ*ExoTUY* triple-mutant strain retained the observed E3 ligase activity (**Fig. 2E**). Interestingly, to varying degrees, the ligase activity is also widely observed among a panel of 14 other *P. aeruginosa* clinical isolates (**Fig. 2F, S2B-C**). These data suggest the presence of an unidentified secreted E3 ligase among the repertoire of ExsA-dependent *P. aeruginosa* virulence factors.

**Figure 2:**
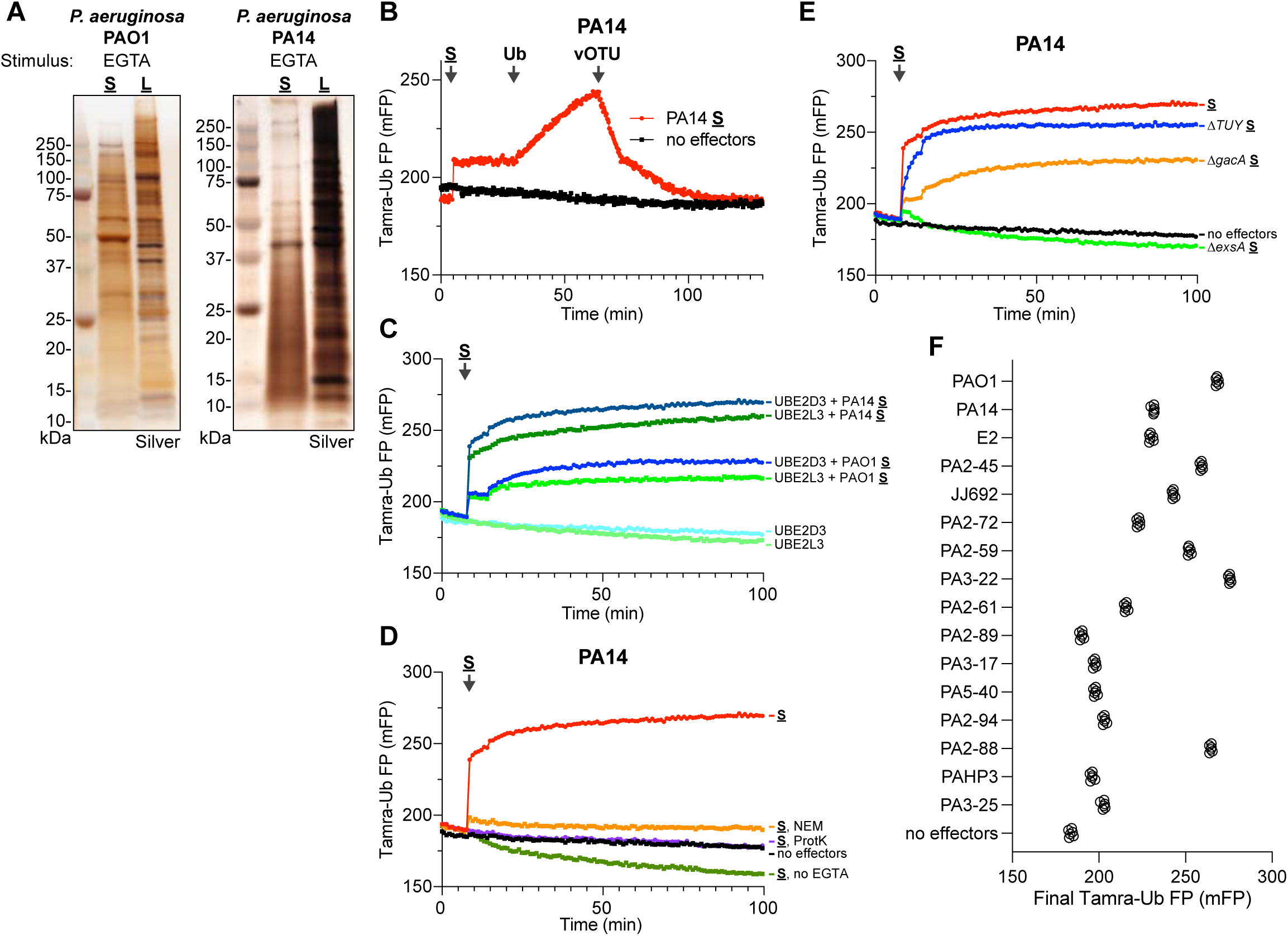
Detection of E3 ligase activity secreted by *P. aeruginosa*. A. Silver-stained SDS-PAGE analysis of secreted (**<u>S</u>**) and lysate (**<u>L</u>**) fractions prepared following EGTA stimulation of *P. aeruginosa* PAO1 and PA14 strains. B. Representative FP traces monitoring the Tamra-Ub ligase substrate following addition of the PA14 secreted fraction. At the indicated timepoints, additional unlabeled Ub or the DUB vOTU were added to stimulate product extension or deconjugation, respectively. C. Representative FP traces monitoring the Tamra-Ub ligase substrate following addition of either the PA14 or PAO1 secreted fractions to reactions containing either the promiscuous E2 UBE2D3 or the Cys-specific E2 UBE2L3. D. Representative FP traces monitoring the Tamra-Ub ligase substrate following addition of the PA14 secreted fraction that was untreated, pre-treated with NEM or Proteinase K (ProtK), or generated without stimulation with EGTA. E. Representative FP traces monitoring the Tamra-Ub ligase substrate following addition of secreted fractions generated from the indicated PA14 mutant strains. F. Final FP values of the Tamra-Ub ligase substrate following addition of secreted fractions from the indicated *P. aeruginosa* clinical isolates.

### Identification of a *P. aeruginosa* E3 ligase

In an initial effort to further purify the secreted ligase activity, the PA14 secreted fraction was resubjected to more refined, stepwise ammonium sulfate precipitations (**Fig. 3A**). This process separated defined protein bands in the 20-50% ammonium sulfate range from the much more complex components in the 50-80% ammonium sulfate range. Among these fractions, the 30-40% ammonium sulfate fraction contained the highest amount of ligase activity (**Fig. 3B, S3A**). Given the Cys-based mechanism of the secreted ligase (**Fig. 2C-D**), the 20-30% and 30-40% secreted fractions were subjected to labeling with the E1-E2-E3 cascading activity-based probe, Ub-DHA, which employs a dehydroalanine (DHA) warhead at its C-terminus to covalently capture Ub-conjugating enzymes^48^. As expected, a biotinylated Ub-DHA probe labeled the E1 (UBE1) and E2 (UBE2D3) in the reaction (**Fig. 3C**). Upon addition of *P. aeruginosa* secreted fractions, additional Ub-DHA-reactive bands were identified that increased in intensity between the 20-30% and 30-40% fractions (**Fig. 3C**). Following visualization with a fluorescently-labeled Ub-DHA probe, these reactive bands were excised and analyzed by mass spectrometry.

**Figure 3:**
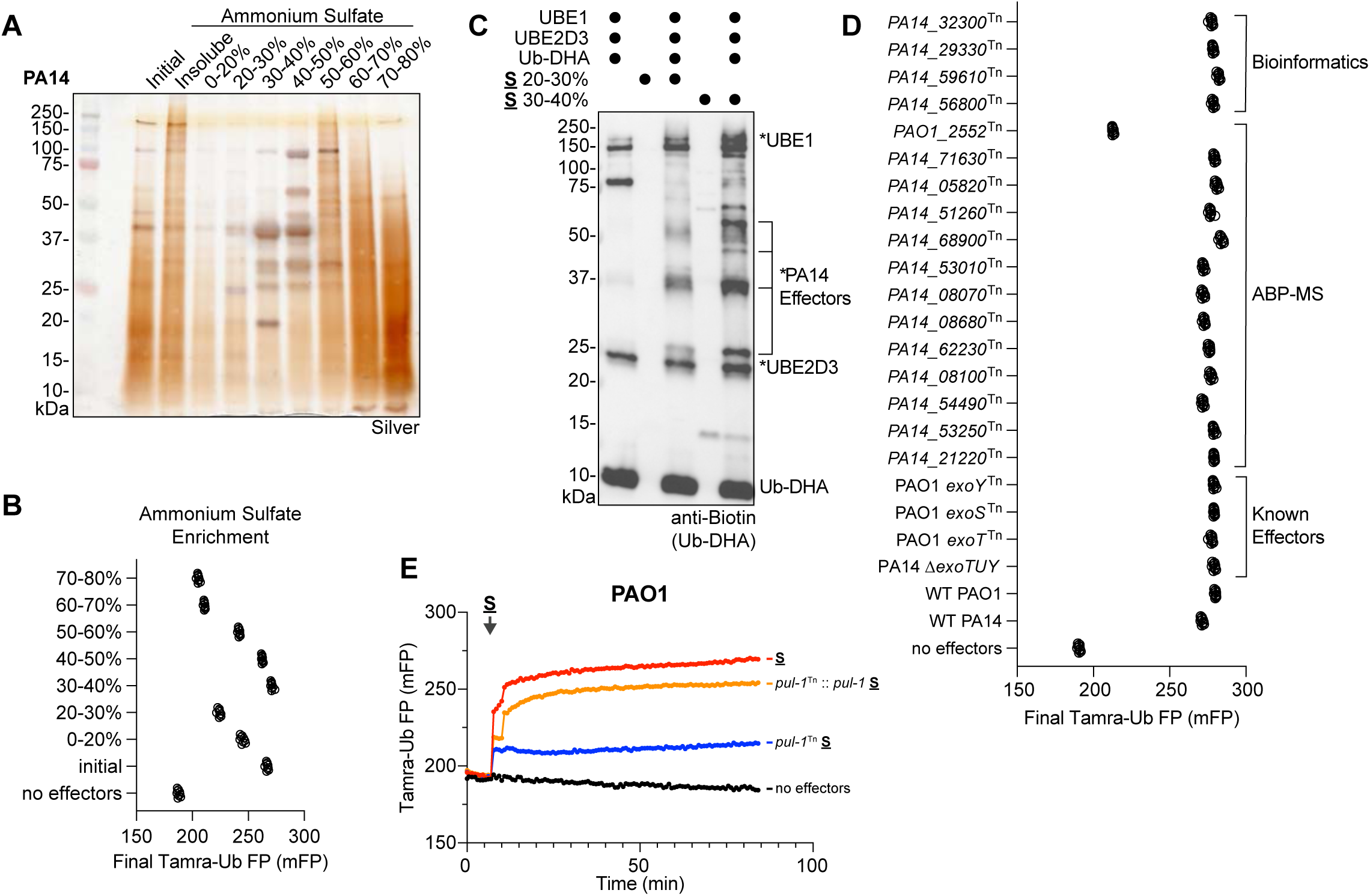
Identification of a *P. aeruginosa* E3 ligase. A. Silver-stained SDS-PAGE analysis of the PA14 secreted fraction following stepwise ammonium sulfate fractionation. B. Final FP values of the Tamra-Ub ligase substrate following addition of the indicated ammonium sulfate fractions of PA14 secreted protein. C. Reactivity of the cascading Ub-DHA activity-based probe with the indicated ammonium sulfate fractions of PA14 secreted protein. Reactions were resolved by SDS-PAGE and visualized by western blot for the biotinylated Ub-DHA probe. Reacted proteins, including putative probe-reactive PA14 effectors, are labeled with an asterisk. D. Final FP values of the Tamra-Ub ligase substrate following addition of secreted fractions generated from the indicated *P. aeruginosa* mutant strains, which test ligase candidates from bioinformatic prediction, ABP-MS analysis, or known effectors. E. Representative FP traces monitoring the Tamra-Ub ligase substrate following addition of secreted fractions generated from PAO1 wild-type, the *pul-1*^Tn^ mutant strain, or a *pul-1*^Tn^ mutant strain complemented with *pul-1*.

Candidate proteins identified from the activity-based probe mass spectrometry (ABP-MS) analysis were combined with candidates identified in a bioinformatics approach using the Ub-SIEVE model^15^, and the resulting list was screened by preparing secreted fractions from the associated PAO1 and PA14 transposon mutant strains for testing in UbiReal^49,50^. Among this list of candidates, as well as the established T3SS effectors, only the *PAO1_2552*^Tn^ strain (carrying an inactivating transposon inserted into PA2552) exhibited a loss in secreted ligase activity (**Fig. 3D**). In a conventional western blot approach, the *PAO1_2552*^Tn^ secreted fraction lacked nearly all ligase activity, compared to the wild-type PAO1 strain (**Fig. S3B**). We therefore renamed PA2552 to *Pseudomonas* Ub ligase 1 (PUL-1). A full kinetic UbiReal profile of the *pul-1*^Tn^ secreted fraction showed a considerable loss in ligase activity, which was restored to wild-type levels when *pul-1* was reintroduced into the transposon mutant strain on a plasmid with its native promoter region (**Fig. 3E**). Under these conditions, PUL-1 therefore appears to be the predominant E3 ligase secreted by *P. aeruginosa*.

### Characterization of PUL-1 E3 ligase activity

To further characterize its ligase activity, the *pul-1* gene was cloned from PAO1 and used for recombinant protein expression in *E. coli* (**Fig. 4A**). An *in vitro* ubiquitination reaction with purified PUL-1 showed robust polyUb chain formation, including unanchored diUb as well as longer, high molecular weight chains (**Fig. 4B**). As was the case with the *P. aeruginosa* secreted fraction, the ligase activity of purified PUL-1 could be reversed with the nonspecific DUB USP21^43^, and additionally could be inhibited by pre-treatment with NEM (**Fig. 4B**). As expected, purified PUL-1 is also labeled with the Ub-DHA probe in a time-dependent manner, and produces an ∼55 kDa band that matches a reactive band observed in the *P. aeruginosa* secreted fraction (**Fig. 4C**). Across a panel of E2 enzymes, PUL-1 is active with the UBE2D family (particularly UBE2D1 and UBE2D2), UBE2L3, and UBE2W (**Fig. 4D, S4A**). The E3-independent activities of UBE2K and UBE2S were also slightly higher in the presence of PUL-1. The form of polyUb produced by PUL-1 was examined using a Ub chain restriction (UbiCRest) analysis, which involves subjecting PUL-1 polyUb products to a panel of linkage-specific DUBs and examining their cleavage patterns^43^. Only treatment with the nonspecific DUBs vOTU and USP21, or the combination of all linkage-specific DUBs, resulted in appreciable cleavage of PUL-1 products (**Fig. 4E**). This behavior is consistent with products comprised of monoubiquitination and nonspecific polyubiquitination. The same behavior is observed with panels of K-only or K-to-R Ub mutants, which indicate no specificity in the type of polyUb formed by PUL-1 (**Fig. S4B-C**). To test whether PUL-1 products are defined by a conventional, Lys-Ub linkage, the conditions of the PUL-1 ubiquitination reaction were optimized to capture the transient, PUL-1∼Ub intermediate, in which Ub is loaded onto the PUL-1 active site Cys (**Fig. 4F**). As expected, this activated PUL-1∼Ub thioester intermediate is reactive toward the reducing agent dithiothreitol (DTT) as well as the amino acid Cys. Among the panel of other amino acids tested, however, PUL-1 was only capable of discharging Ub onto Lys, suggesting a specificity toward conventional Ub linkages (**Fig. 4F**). Consistently, mass spectrometry analysis of an *in vitro* PUL-1 ubiquitination reaction identified three sites of PUL-1 Lys auto-ubiquitination as well as formation of K6, K11, K27, K33, K48, and K63 polyUb linkages (**Fig. S4D**).

**Figure 4:**
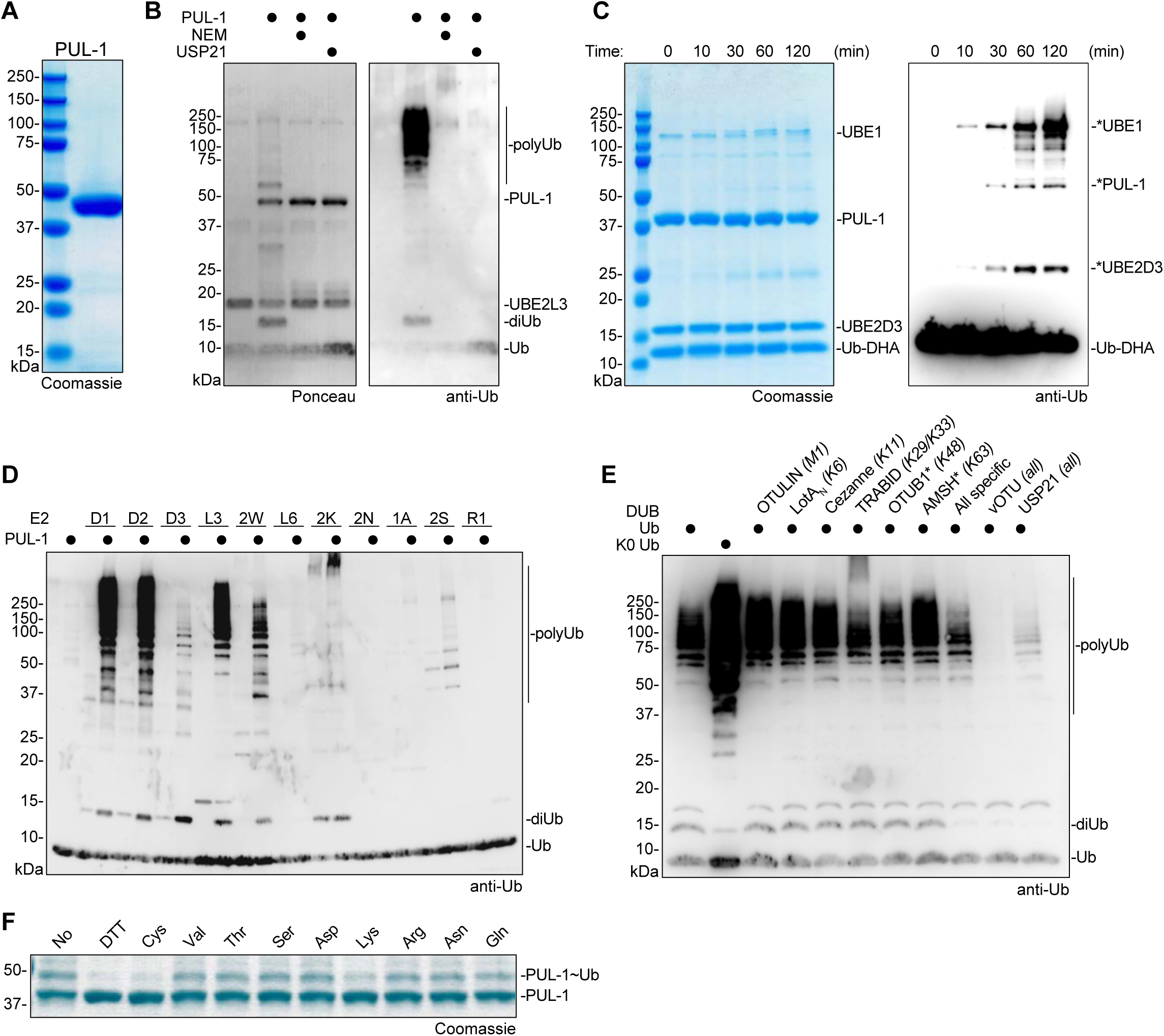
Characterization of PUL-1 E3 ligase activity. A. Coomassie-stained SDS-PAGE analysis of recombinantly purified PUL-1. B. E3 ligase assays for recombinant PUL-1, including conditions with pre-treatment of NEM or post-treatment of the DUB USP21. Reactions were resolved by SDS-PAGE and visualized by Ponceau stain and anti-Ub western blot. C. Time course monitoring reactivity of the cascading Ub-DHA activity-based probe with recombinant PUL-1. Reactions were resolved by SDS-PAGE and visualized by Coomassie stain or anti-Ub western blot. Reacted proteins are labeled with an asterisk. D. E3 ligase assays for recombinant PUL-1 and the indicated panel of E2 enzymes. Reactions were resolved by SDS-PAGE and visualized by anti-Ub western blot. E. UbiCRest assay of PUL-1 ligase products. A PUL-1 ligase reaction was treated with the indicated panel of linkage-specific DUBs, a combination of all linkage-specific DUBs, or the indicated nonspecific DUBs. Reactions were resolved by SDS-PAGE and visualized by anti-Ub western blot. F. Amino acid reactivity analysis of the activated PUL-1∼Ub thioester intermediate. PUL-1 was loaded with Ub, and discharge was monitored following addition of DTT or the indicated amino acids. Reactions were resolved by SDS-PAGE and visualized by Coomassie stain.

### Structural analysis of the PUL-1 E3 ligase fold

Although no experimental structure of PUL-1 is currently available, the AlphaFold2 model exhibits very high confidence, with predicted local distance difference test (pLDDT) scores of >90 throughout most of the 375-residue sequence^51^ (**Fig. 5A**). The modeled structure shows high similarity to an acyl-CoA dehydrogenase fold, and a Dali structural homology analysis highlights high similarity to short chain acyl-CoA dehydrogenases from bacterial as well as eukaryotic origin^52^. Crystal structures of homologous folds from *Burkholderia thailandensis* BTH_II1803 and the rat mitochondrial short-chain specific acyl-CoA dehydrogenase (SCAD) align to the PUL-1 model with less than 1 Å RMSD^53^ (**Fig. 5B**). Residues at the acyl-CoA-binding site, as well as the cofactor FAD-binding site, are highly conserved among PUL-1 and rat SCAD (**Fig. S5A-B**). Despite this similarity, PUL-1 demonstrated no dehydrogenase activity against an octanoyl-CoA substrate, unlike the highly similar *Mycobacterium tuberculosis* FadE13 enzyme (**Fig. 5C**). Conversely, unlike PUL-1, *M. tuberculosis* FadE13 demonstrated no E3 ligase activity (**Fig. 5D**). Located at the putative acyl-CoA-binding site, a PUL-1 E361A mutant had no effect on either the lack of dehydrogenase activity or the apparent E3 ligase activity (**Fig. 5C-D**).

**Figure 5:**
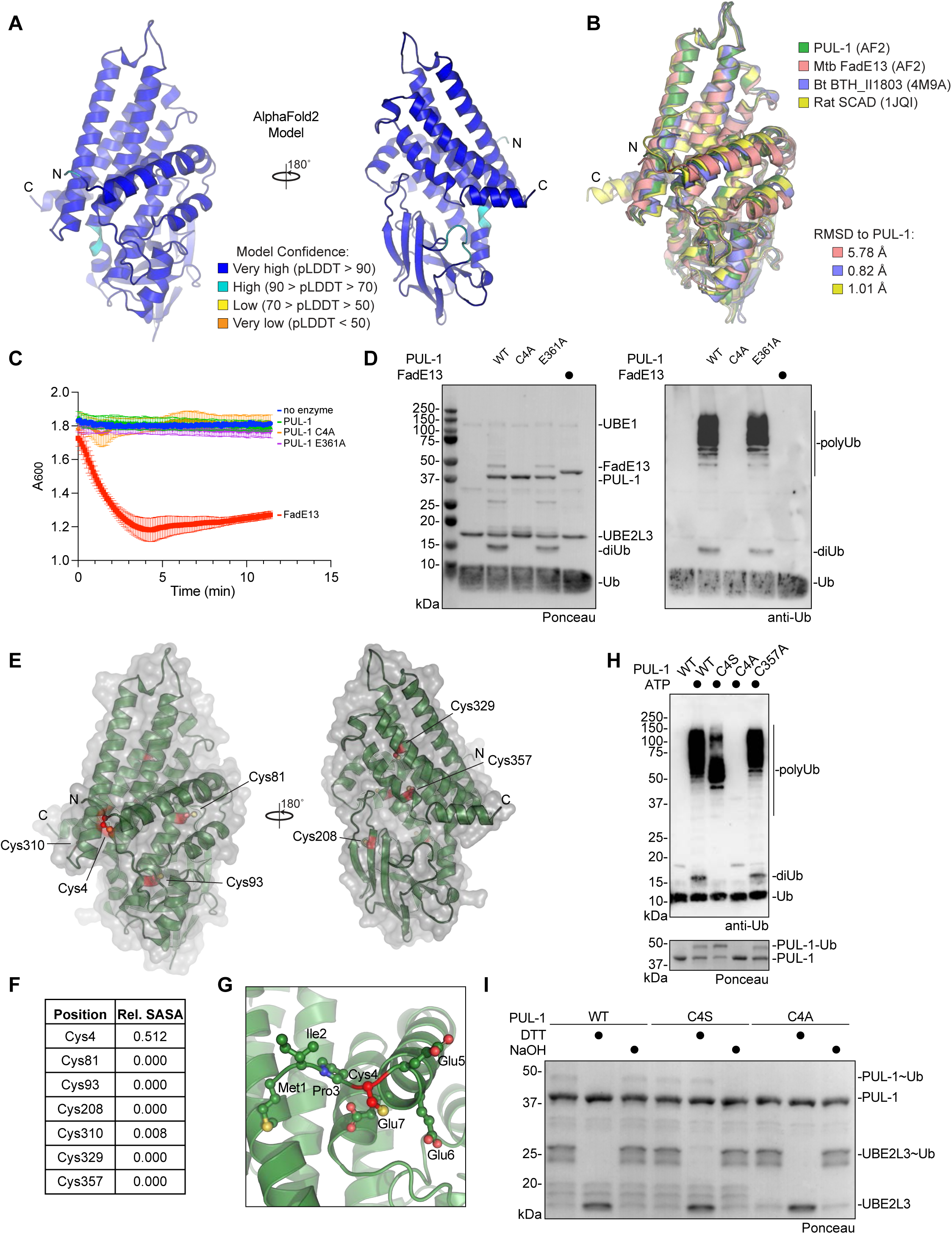
Structural analysis of the PUL-1 ligase fold. A. AlphaFold2 model of PUL-1, colored by pLDDT confidence scores, with N- and C-termini labeled. B. Structural overlay of the AlphaFold2 models of PUL-1, putative acyl-CoA dehydrogenases from *M. tuberculosis* (Mtb) and *B. thailandensis* (Bt, PDB: 4M9A), and the mitochondrial short-chain specific acyl-CoA dehydrogenase (SCAD, PDB: 1JQI) from rat. C-alpha root mean square deviation (RMSD) values are listed. C. Octanoyl-CoA dehydrogenase activity assay monitoring reduction of DCPIP at 600 nm wavelength. D. E3 ligase assays for recombinant PUL-1 and FadE13. Reactions were resolved by SDS-PAGE and visualized by Ponceau stain and anti-Ub western blot. E. AlphaFold2 model of PUL-1 (green), with surface representation (grey) and all Cys residues highlighted (red). F. Relative solvent-accessible surface area (SASA) of each Cys residue within the PUL-1 AlphaFold2 model. G. Detailed view of Cys4 within the PUL-1 AlphaFold2 model, with Cys4 (red) and neighboring residues shown as ball-and-stick. H. E3 ligase assays for wild-type or the indicated PUL-1 mutants. Reactions were resolved by SDS-PAGE and visualized by Ponceau stain and anti-Ub western blot. I. Chemical stability toward reducing agent (DTT) or basic pH (NaOH) of the activated UBE2L3∼Ub and PUL-1∼Ub intermediates. Enzymes were loaded with Ub, and discharge was monitored following the indicated chemical treatments. Reactions were resolved by SDS-PAGE and visualized by Ponceau stain.

As PUL-1 represents an unprecedented E3 ligase fold, we turned toward interpreting the AlphaFold2 model in this perspective. Biochemical evidence suggested a Cys-based ligase mechanism (**Fig. 4B-D**). Analysis of the relative surface accessible surface areas (SASA) for all PUL-1 Cys residues highlighted Cys4 as the only one at the protein surface that could serve as an active site (**Fig. 5E-F**). Located near the N-terminus, Cys4 directly precedes the first α-helix of the PUL-1 fold (**Fig. 5G**). Mutation of Cys4 to alanine ablated PUL-1 ligase activity, whereas an analogous mutation at Cys357 had no effect (**Fig. 5H**). Consistent with trapping the PUL-1∼Ub intermediate as a more stable oxyester linkage, mutation of Cys4 to serine resulted in a lower molecular weight auto-ubiquitination product and no formation of unanchored diUb (**Fig. 5H**). To confirm this prediction, we returned to assay conditions that allow observation of the early PUL-1∼Ub intermediate. While the wild-type PUL-1∼Ub thioester intermediate was susceptible to reduction with DTT, the intermediate formed with the C4S mutant was not, but instead could be hydrolyzed by base treatment (**Fig. 5I**). Meanwhile, the PUL-1 C4A mutant showed no PUL-1∼Ub formation, confirming Cys4 as the active site. Lastly, the C4A mutation had no effect on the lack of dehydrogenase activity demonstrated by PUL-1 (**Fig. 5C**). Among the tested *P. aeruginosa* clinical isolates with secreted ligase activity (**Fig. 2F**), the PUL-1 sequence is highly conserved and all examples carry the catalytic Cys4 residue (**Fig. S5C**). Interestingly, sequence analysis outside of *P. aeruginosa* also identifies related acyl-CoA dehydrogenases with Cys or Ser residues at this active site position, suggesting a wider adaptation of E3 ligase activity into this common protein fold (**Fig. S5D**).

### PUL-1 ligase activity modulates *P. aeruginosa* virulence

As *P. aeruginosa* PUL-1 could only function as an E3 ligase within the environment of a eukaryotic host where the E1, E2, and Ub are present, we sought to assess its role as a virulence factor. Consistent with this role, PUL-1 had no impact on the doubling time of *P. aeruginosa* in culture (**Fig. 6A**). Furthermore, PUL-1 had no effect on *P. aeruginosa* motility, measured either through swimming or swarming (**Fig. S6A-D**). Overall, the mechanisms of *P. aeruginosa* pathogenesis are broadly conserved across mammalian, plant, and metazoan hosts. To determine the role of PUL-1 in virulence, we used an established *C. elegans* model system^54^. *P. aeruginosa* infects and kills *C. elegans* through a process that mirrors infection in other hosts and correlates with bacterial accumulation in the intestine.

**Figure 6:**
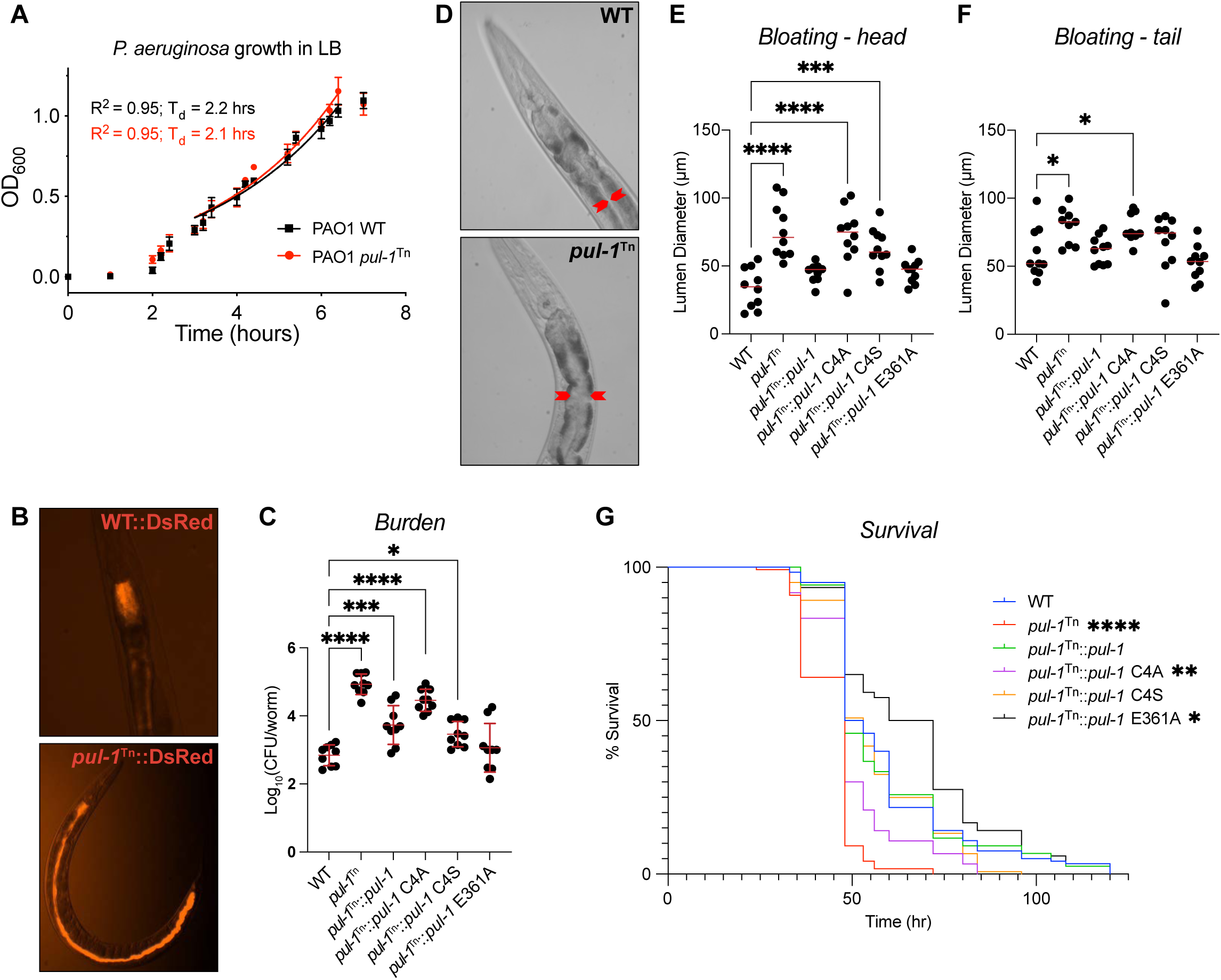
PUL-1 ligase activity modulates *P. aeruginosa* virulence. A. Growth curves for wild-type (WT) and the PAO1 *pul-1*^Tn^ mutant strain in LB, with doubling times (T_d_) indicated. B. Representative fluorescent images of *C. elegans* infected with WT and the PAO1 *pul-1*^Tn^ mutant strain expressing DsRed. C. Bacterial burden measured as colony-forming units (CFU) per worm, following infection with the indicated PAO1 strains. Mean values and standard deviation are indicated in red. Significance was determined by one-way ANOVA with Dunnett’s multiple comparisons test. D. Representative images of *C. elegans* intestinal bloating near the head, following infection with WT or the PAO1 *pul-1*^Tn^ mutant strain. The intestinal lumen diameter is indicated by red arrows. E. Quantification of *C. elegans* intestinal lumen diameter near the head following infection with the indicated PAO1 strains. Median values are shown in red. Significance was determined by one-way ANOVA with Dunnett’s multiple comparisons test. F. As in **E)**, for intestinal lumen diameters near the tail. G. Survival curves for *C. elegans* infected with the indicated PAO1 strains. Significance was determined by the Log-rank (Mantel-Cox) test.

To visualize the extent of intestinal infection, *P. aeruginosa* strains expressing DsRed fluorescent protein were used to infect worms prior to visualization by fluorescence microscopy. While the wild-type *P. aeruginosa* infection remained localized near the worm pharynx under these conditions, in stark contrast the *pul-1*^Tn^ strain occupied the entire length of the intestine (**Fig. 6B**). Accordingly, quantification of the bacterial burden by colony-forming units (CFU) revealed a several log-fold increase following infection with the *pul-1*^Tn^ strain, compared to wild-type (**Fig. 6C**). This effect on bacterial burden could be complemented with a wild-type *pul-1* transgene expressed on a plasmid under its native promoter region (**Fig. 6C**). To test whether this impact on virulence was linked to PUL-1 ligase activity, the *pul-1*^Tn^ mutant strain was instead complemented with the structure-guided PUL-1 mutants. Complementation with the C4A mutation, which ablated PUL-1 ligase activity (**Fig. 5H**), mimicked the *pul-1*^Tn^ mutant strain and exhibited a higher bacterial burden (**Fig. 6C**). Interestingly, the C4S mutant, which only restricted PUL-1 ligase activity (**Fig. 5H**), displayed an intermediate phenotype, whereas the E361A mutant at the putative acyl-CoA-binding site behaved like wild-type (**Fig. 6C**).

Consistent with a higher bacterial burden, nematodes infected with the *pul-1*^Tn^ strain exhibited a dramatic intestinal bloating phenotype compared to those fed with wild-type *P. aeruginosa* (**Fig. 6D**). This bloating phenotype was observed at both the head and the tail of the worms, and could be restored to wild-type levels following complementation of the *pul-1*^Tn^ mutant strain with a *pul-1* transgene (**Fig. 6E-F, S6E**). Complementation with the C4A and C4S mutations in the PUL-1 ligase active site resulted in significant intestinal bloating similar to the *pul-1*^Tn^ strain (**Fig. 6E-F, S6E**). Meanwhile, complementation with the E361A mutation at the putative acyl-CoA-binding site restored bloating to levels observed with wild-type *P. aeruginosa* (**Fig. 6E-F, S6E**).

The elevated burden and bloating phenotypes were associated with a significant decrease in the lifespan of worms infected with the *pul-1*^Tn^ strain, compared to wild-type (**Fig. 6G**). Once again, this effect was dependent upon the ligase activity of PUL-1, as infection with the C4A-complemented mutant strain caused a similar reduction in lifespan to the *pul-1*^Tn^ strain (**Fig. 6G**). Surprisingly, infection with the C4S-complemented mutant strain resulted in a lifespan similar to the wild-type *P. aeruginosa* infection, suggesting that the restricted ligase activity of the serine substitution retains some level of biologically relevant function. Infection with the E361A-complemented strain, meanwhile, did not decrease lifespan compared to the wild-type infection. Altogether, the *C. elegans* infections demonstrate a ligase-dependent role of PUL-1 in regulating the virulence of *P. aeruginosa*.

## DISCUSSION

The discovery of PUL-1 as a cryptic E3 Ub ligase lends further weight to the evolutionary advantage of regulating Ub signaling during bacterial infection. That PUL-1 exhibited more of an antivirulence role under these infection conditions was surprising, but not without precedent^55,56^. Other infection models may reveal a more traditional role, or PUL-1 might function to modulate the level of *P. aeruginosa* virulence. While many bacterial ligases direct Ub-dependent degradation of their targets, others instigate nondegradative signaling^10^. The identity and fate of PUL-1 substrates, and their ties to its role in modulating virulence, will be an interesting area of future research. One potential clue is the observation that a PUL-1 C4S mutant complemented many of the phenotypes observed during *C. elegans* infection with the *pul-1*^Tn^ strain. Based on our knowledge of eukaryotic Cys-based ligases, a serine mutation in the active site should stabilize the E3∼Ub intermediate and reduce or eliminate subsequent transfer. This is consistent with the restricted ligase activity of the PUL-1 C4S mutant observed *in vitro*. That the C4S mutant functions similarly to wild-type PUL-1 *in vivo*, however, suggests that either transfer from a serine active site is possible under these conditions, or that the role of PUL-1 is to ubiquitinate itself. Interestingly, many PUL-1 orthologues in other bacteria encode a serine at this catalytic position, suggesting that some form of Ub ligase activity may be retained more broadly.

The finding that PUL-1 catalyzes Ub conjugation through an acyl-CoA dehydrogenase protein fold illustrates the importance of an unbiased approach in studying bacterial Ub regulators. PUL-1 is not alone in this regard either; the SidE ligase family from *L. pneumophila* combines mono-ADP-ribosyltransferase and phosphodiesterase activities to catalyze noncanonical, ATP-independent ubiquitination^57^. While our work has identified the catalytic center for PUL-1 ligase function, how it engages with eukaryotic E2s for Ub transfer remains a topic for future work. Structurally, PUL-1’s ligase activity is spatially distinct from its putative acyl-CoA dehydrogenase function. Though we do not observe dehydrogenase activity *in vitro*, we cannot exclude that PUL-1 retains moonlighting activities. Moonlighting functions are not uncommon among viral and bacterial proteins, even in conjunction with Ub regulation. Bacterial DUBs from the CE clan of cysteine proteases can exhibit mixed Ub/Ub-like protease as well as acetyltransferase activities through the same catalytic center^33,58^. In the case of PUL-1, however, the spatially removed Cys4 active site allowed for targeted elimination of specifically its ligase function, thereby revealing an important role of PUL-1 ligase activity *in vivo*.

The approach we have developed to identify natively secreted Ub regulators offers several key advantages over previous methods. First, it eliminates several biases that hinder the identification of convergent activities: a) it is agnostic toward any sequence or structural similarities to known Ub regulators, b) it avoids any presumptions about the mechanisms of Ub regulation, such that they would react with activity-based probes, and c) it allows for discovery of previously uncharacterized virulence factors. Secondly, the approach is highly versatile, both in the input sample of bacterial proteins as well as in the reaction components. Through pre-treatment of secreted fractions, or through swapping out the E2 in ligase assays, we could learn a great deal about the nature of Ub regulation, before even identifying the responsible enzyme. It would also be straightforward to introduce E2 or Ub mutations into the assay, in order to test dependency on common interaction surfaces. Furthermore, though we focused on Ub, one could test for conjugating, deconjugating, or binding activities toward ubiquitin-like modifiers by simply exchanging the enzymes and substrates. A third key advantage of this approach is its synergy with other methods. After observing and studying the activity in the mixture of secreted proteins, one can quickly select a suitable activity-based probe for capture, enrichment, and identification of the responsible enzyme. Alternatively, one could further purify the activity through biochemical properties and fractionation, or through noncovalent interaction with the reaction components. Lastly, where available, transposon mutant libraries offer a streamlined approach to testing candidate proteins. Highlighted by benchmark studies in three bacteria and the successful identification of a cryptic *P. aeruginosa* E3 ligase, this approach offers a compelling opportunity to identify Ub regulators that modulate host-pathogen interactions and disease.

Though our approach offers many benefits in terms of eliminating bias and incorporating versatility, there are several limitations in its current form. One limitation is the method used to generate and harvest secreted effectors. While the stimuli we selected are widely utilized, they are most likely not perfect mimics of a host infection and therefore may not trigger secretion to the same extent. An alternative, which we took advantage of in the case of *S. flexneri*, is to also test a mutant strain that constitutively secretes even in the absence of stimulation. Another limitation is the lack of a eukaryotic environment in our assays for Ub regulation. This could mean that cofactors or substrates required for activity are missing. While some effectors require eukaryotic cofactors for function^21,55,59^, this has not been observed for Ub regulation, likely because such an activity would not be toxic to the bacterium prior to secretion and thus there would be no selective pressure to establish spatiotemporal control. Incorporating protein-depleted eukaryotic lysates into the approach could also account for required cofactors. Regarding substrates, the monoUb-based substrate we used for monitoring DUB activity is broadly applicable, but to detect linkage-specific DUB activities, fluorescent polyUb substrates could be used instead^60^. As for ligase activities, most E3 ligases will auto-ubiquitinate or produce unanchored polyUb chains, even in the absence of substrate. Our initial approach for detecting ligase activity also accounted for E2 selectivity by incorporating highly promiscuous E2s, but additional E2s could easily be tested in parallel. Thus, with minor modifications, our approach can be broadly applicable to diverse bacteria and forms of Ub regulation.

## MATERIALS AND METHODS

### Bacterial strains and growth conditions

*S.* Typhimurium SL1344 and secretion mutant strains were a kind gift from Dr. Leigh Knodler (U. Vermont). EPEC O127:H6 strain E2348/69 was a kind gift from Dr. Brett Finlay (UBC). *S. flexneri* M90T and secretion mutant strains were a kind gift from Dr. John Rohde (Dalhousie U.). *P. aeruginosa* PAO1 and PA14, along with associated transposon or deletion mutant strains, were accessed from the Ausubel and Manoil strain libraries^49,50^. *P. aeruginosa* isolate PAHP3 was used previously^61^, and isolates JJ692 and E2 are part of a 20-strain diversity panel used previously^62^. Other *P. aeruginosa* clinical isolates are from a panel of multidrug-resistant strains isolated from pediatric patients with cystic fibrosis, described previously^63^. *S. flexneri* strains were cultured in Trypsic Soy Broth (TSB). All other strains were cultured in Luria-Bertani broth (LB) at 37 °C with the following antibiotics, as required: Gentamicin 15 µg/mL, Tetracycline 5 µg/mL, Carbenicillin 200 µg/mL, Streptomycin 50 µg/mL, and Kanamycin 50 µg/mL.

### Cloning and mutagenesis

The *PA2552* gene of *P. aeruginosa* was PCR amplified from PAO1 genomic DNA using primers PA2552F *(5’-* AAGTTCTGTTTCAGGGCCCGatgattccctgcgaagaagag*-3’)* and PA2552R *(5’-* ATGGTCTAGAAAGCTTTActacaggctgcgcacg*-3’)* or PA2552F_pUCP18 (5′-tcagatGGATCCcaacgtccacggcg-3′) and PA2552R_pUCP18 (5′-tgagatAAGCTTctacaggctgcgcg-3′). The amplified PA2552F/R DNA fragment was combined 1:3 vector:insert with linearized pOPIN-B and transformed without ligation into TOP10 *E. coli* utilizing a rec-independent process. The amplified PA2552F/R_pUCP18 DNA fragment was digested with BamHI and HindIII before ligation into pUCP18 and propagation in TOP10 *E. coli.* PA2552 point mutations C4A, C357A, and E361A of were performed using Quikchange PCR with PA2552_pUCP18 or PA2552_pOPINB as template and primers PA2552C4AF (*5’-* TCCCgcgGAAGAAGAGATCCAGATCCGT*-3’*) and PA2552C4AR (*5’-* CTTCcgcGGGAATCATCGCGGGT*-3’*), or PA2552C4AF (*5’-* TCCCtcaGAAGAAGAGATCCAGATCCGT*-3’*) and PA2552C4AR (*5’-* CTTCtgaGGGAATCATCGCGGGT*-3’*), or PA2552E361AF (*5’-* CTACgcgGGCAccagcgacgt*-3’*) and PA2552E361R (*5’-* TGCCcgcGTAGatctggcagaccc*-3’*) or PA2552C357F (*5’-GGTCgcgCAGATCTACGAGGG-3’*) and PA2552C357R (*5’-TCTGcgcGACCCGCACGTCCC-3’*). Electrocompetent *P. aeruginosa* were prepared using the sucrose method according to Choi et al^64^. Transformants carrying the pUCP18 complementation plasmid were selected on LB agar with Carbenicillin (200 µg/mL).

### Collection of secreted effectors

Bacteria were plated onto agar containing selective antibiotics and grown overnight at 37 °C. All effector preparations were generated from 25-50 mL of starting culture. For strains of *P. aeruginosa,* single colonies were inoculated into sterile LB containing 5 mM EGTA and grown at 37 °C with shaking at 215 rpm overnight. To harvest *S.* Typhimurium SPI-I effectors, strains were first grown overnight in LB at 37 °C, then diluted from the overnight culture 1:300 in fresh LB containing 300 mM NaCl and allowed to grow for 3 hours to mid-log phase at 37 °C with shaking at 215 rpm. For isolation of *S.* Typhimurium SPI-II effectors, overnight cultures were diluted 1:30 in MgM-MES media containing 170 mM 2-[N-morpholino]ethane-sulfonic acid (MES), 5 mM KCl, 7.5 mM (NH_4_)_2_SO_4_, 0.5 mM K_2_SO_4_, 1 mM KH_2_PO_4_, 8 µM MgCl_2_, 38 mM glycerol, and 0.1% casamino acids, set to a final pH of 5.0. The cultures were grown at 37 °C for 4 hours with shaking at 215 rpm, then the cells were pelleted and resuspended in 1 mL of 25 mM sodium phosphate (pH 7.4), 150 mM NaCl and left standing at 37 °C for 1 hour. Red colonies of *S. flexneri*, observed on TSB agar plates containing Congo red, were grown in liquid TSB overnight at 37 °C with shaking at 215 rpm. Overnight cultures were then diluted 1:300 in TSB and allowed to grow for 4 hours to mid-log phase at 37 °C with shaking at 215 rpm. Strains of EPEC were grown overnight at 37 °C with shaking at 215 rpm in LB. Bacteria were then sub-cultured 1:40 in pre-warmed Dulbecco’s modified eagle medium (DMEM) and allowed to grow standing at 37 °C with 5% CO_2_ for 6 hours (until OD_600_ reached 1).

Following stimulation of effector secretion, bacteria were pelleted by centrifugation at 2,400 xg for 25 min. Pellets were freeze-thawed twice and treated with protease inhibitor cocktail (MilliPore-Sigma) and 1 µg/mL lysozyme in 1 mL of 25 mM sodium phosphate (pH 7.4), 150 mM NaCl, 1 mM DTT for 30 min on ice. The lysate was clarified by centrifugation (29,000 xg, 10 min) and sterile filtering. Culture supernatants were sterile filtered and Tris buffer (pH 8.0) was added to a final concentration of 25 mM before slowly adding ammonium sulfate powder to a final percentage of 75% w/v. Ammonium sulfate was allowed to dissolve at 4 °C for 30 min with light stirring. Precipitated proteins were collected by centrifugation at 35,000 xg for 30 min. Pellets were resuspended in 0.5-1 mL of 25 mM sodium phosphate (pH 7.4), 150 mM NaCl, 0.5 mM DTT and dialyzed against the same buffer to remove excess ammonium sulfate. Samples were subjected to Bradford assay to determine total protein concentration, normalized to 1-2 mg/mL, flash frozen in liquid nitrogen, and stored at -80 °C.

### SDS-PAGE analysis of secreted effector pools

Samples of secreted and lysate fractions (5 ug) from each pathogen were diluted in reducing SDS sample buffer and boiled for 5 mins at 98 °C. Proteins were resolved by 4-12% Tris-Glycine SDS-PAGE (BioRad). Gels were fixed and silver stained according to the manufacturer’s protocol (BioRad).

### Recombinant protein production

All recombinant proteins were produced in Rosetta *E. coli* (Millipore). Transformed cells were cultured in LB at 37 °C until an OD_600_ of 0.6-0.8, at which point protein expression was induced with 0.2 mM IPTG and growth continued at 18 °C overnight. Cells were harvested by centrifugation at 4,000 xg and resuspended in Buffer A: 25 mM Tris (pH 7.4), 200 mM NaCl, 2 mM 2-mercaptoethanol (with the exception of UBE1, which lacked reducing agent). Following a freeze-thaw, cells were treated with protease inhibitor cocktail (Millipore-Sigma), 50 µg/mL PMSF, 50 µg/mL DNase, and 200 µg/mL lysozyme for 30 min on ice. Samples were then lysed by sonication and clarified by centrifugation at 35,000 xg for 30 min. UBE1was purified by activation onto GST-Ub-loaded glutathione resin, washed with 25 mM Tris (pH 7.4), 200 mM NaCl, and eluted with the same buffer containing 10 mM DTT. The resulting elution was further purified by size exclusion chromatography (Superdex75 pg, Cytiva). UBE2D3 and UBE2L3 were expressed without affinity tags and purified by cation exchange of clarified lysate in 30 mM MES pH 6.0, 1 mM EDTA, followed by size exclusion chromatography in 25 mM sodium phosphate pH 7.4, 150 mM NaCl (Superdex75 pg, Cytiva).

His-tagged PUL-1 constructs were expressed as above and purified with cobalt resin using standard procedures (ThermoFisher). Proteins were eluted with Buffer A containing 300 mM imidazole and subjected to size exclusion chromatography in 50 mM HEPES (pH 8), 150 mM NaCl, 0.5 mM DTT (Superdex75 pg, Cytiva). All proteins were concentrated using Amicon centrifugal filters, quantified by absorbance, and flash frozen for storage at -80 °C.

### Fluorescence polarization assays

FP DUB assays were adapted from Pruneda et al^65^. A master solution with a final concentration of 25 mM Tris (pH 7.4), 100 mM NaCl, 5 mM 2-mercaptoethanol, 0.1 mg/mL BSA, and 100 nM Ub-KG(Tamra) was added to sample wells. FP measurements were made at room temperature using a BMG LabTech ClarioStar with an excitation wavelength of 540 nm, an LP 566 nm dichroic mirror, and an emission wavelength of 590 nm. FP was monitored for 10 cycles before adding 10 µg of secreted or lysate fractions from *S.* Typhimurium and allowing the reaction to continue for 2.5 hours. Each sample was prepared in triplicate, with the FP values averaged over time points.

FP ligase assays were adapted from Franklin et al^36,37^. A master solution with a final concentration of 25 mM sodium phosphate (pH 7.4), 150 mM NaCl, 10 mM MgCl_2_, 100 nM N-terminally labeled Tamra-Ub, 125 nM UBE1, 2 µM E2 (UBE2D3 or UBE2L3) was added to sample wells. To generate E2∼T-Ub conjugate, 1 µl of 100 mM ATP was added to the appropriate sample wells (5 mM final). FP measurements were made at room temperature using a BMG LabTech ClarioStar with an excitation wavelength of 540 nm, an LP 566 nm dichroic mirror, and an emission wavelength of 590 nm. FP was monitored for 10 cycles before adding 10 µg of secreted or lysate fractions from a given pathogen. Secreted fractions that were previously boiled for 10 min at 98 °C, pre-treated with Proteinase K, or pre-treated with NEM, were used as negative controls for ligase activity. FP was monitored for another 10 cycles to visualize early Ub transfer events and/or noncovalent interactions between bacterial proteins and Tamra-Ub. Subsequently, excess unlabeled Ub was introduced by adding 3 µl of 250 µM Ub, for a final concentration of 37.5 µM. FP was then monitored for an additional 2 hours. Each sample was prepared in triplicate, with the FP values averaged over time points.

### Gel-based ubiquitination assays

10 µg of lysate/effector pool or 2 µM of purified PUL-1 were mixed with 125 nM E1, 2 µM E2, 37.5 µM Ub, 10 mM MgCl_2_, and 5 mM ATP in 25 mM sodium phosphate (pH 7.4), 150 mM, NaCl, 0.5 mM dithiothreitol (DTT). The reactions were incubated at 37 °C for 80 min and terminated by the addition of reducing SDS sample buffer.

### Preparation of biotin-Ub-DHA

Preparation of the Ub-DHA probe was adapted from Mulder et al^48^. Ubiquitin with an N-terminal AviTag and G76C mutation was incorporated into pOPIN-B. Protein expression and cell lysis were performed as described above. The resulting clarified lysate was passed over pre-equilibrated cobalt resin, allowing his-tagged Ub to bind (ThermoFisher). Resin was washed with 1 L of Buffer A before eluting with Buffer A containing 300 mM imidazole. Eluted protein was then concentrated with Amicon centrifugal filters and subjected to a biotinylation reaction consisting of 100 µM Avi-tagged protein, 5 µl of 1M MgCl_2_, 20 µl of 100 mM ATP, 20 µl of 50 µM GST-BirA, and 3 µl of 50 mM D-Biotin in 1 mL of 25 mM sodium phosphate (pH 7.4), 150 mM NaCl. The biotinylation reaction proceeded for 1 hr at 30 °C with light shaking. Following size exclusion chromatography into of 25 mM sodium phosphate (pH 7.4), 150 mM NaCl (Superdex75 pg, Cytiva), the resulting biotin-Ub-Cys product was treated with 5 mM DTT to reduce the C-terminal cysteine, desalted into 50 mM sodium phosphate (pH 8.0), and immediately reacted with 500-fold excess dibromohexandiamide (prepared at 300 mM in DMSO). The reaction was incubated at room temperature for 3 hours, and excess dibromohexandiamide was removed by desalting.

### Mass spectrometry sample preparation, in-gel digestion, and LC-MS

To facilitate visualization and excision of Ub-DHA-reactive bands within the *P. aeruginosa* effector pool, a fluorescent Cy5-labled Ub-DHA probe was generated by total synthesis^48^. 30 µg of Cy5-Ub-DHA was incubated with 0.5 mg of *P. aeruginosa* effector pool, 0.5 µM E1, and 5 µM UBE2D3 along with 10 mM MgCl_2_ and 10 mM ATP for 2 hours at 37 °C. Reactions were run into 4-12% SDS-PAGE gels, scanned at 658 nm for Cy5, and Coomassie stained. Gel bands corresponding to 30, 40, and 50 kDa were excised from Ub-DHA-reacted and control samples, sliced into small pieces, and incubated in 500 μl acetonitrile at room temperature for 10 minutes. Acetonitrile was removed, and disulfide bonds were reduced by adding 50 μl of 10 mM DTT in 100 mM ammonium bicarbonate and incubation at 56 °C in a thermomixer with shaking at 400 rpm for 30 minutes. Samples were cooled to room temperature and washed once with 500 μl acetonitrile. After removing acetonitrile, gel slices were destained by addition of 100 μl of 100 mM ammonium bicarbonate/acetonitrile (1:1 v/v) and incubation at room temperature for 30 min in a thermomixer with shaking at 400 rpm. Destaining solution was discarded, and gel slices were incubated in 500 μl acetonitrile for 20 minutes at RT followed by decanting of acetonitrile wash. Proteins were digested by submerging gel slices in 50 μl of freshly prepared sequencing grade trypsin (#22720, Affymetrix, Santa Clara, CA) at 13 ng/μl in 10 mM ammonium bicarbonate containing 10% acetonitrile (v/v). Digestion was performed overnight in a thermomixer at 37 °C with heated lid and shaking at 400 rpm. Peptides were extracted by addition of extraction buffer (1:2 (v/v) 5% formic acid/acetonitrile) at a 2:1 extraction buffer/sample (v/v) ratio, and incubation for 15 min at 37 °C. Peptides were concentrated to dryness in a vacuum centrifuge, resuspended in 10 μl of 5% acetonitrile, and transferred to a new vial. Residual peptides were collected by washing the original tryptic peptide tube with 10 μl water and added to the new vial.

Approximately 500 ng of peptides were injected onto a primary trap column (4 cm x 150 µm i.d. packed with 5 µm Jupiter C18 particles (Phenomenex, Torrence, CA)) and separated in a capillary column (70 cm x 75 µm i.d. packed with 3 µm Jupter C18 particles). Peptides were separated with 0.1% formic acid in acetonitrile (mobile phase B) and 0.1% formic acid in water (mobile phase A) at a 300 nl/min flow rate. The elution gradient consisted of 20 minutes at 12% B, 75 min at 30% B, and 97 min at 45% B. Eluting peptides were analyzed by an inline quadrupole orbitrap mass spectrometer (Q-Exactive HF, Thermo Fisher Scientific, San Jose, CA). Spectra were collected in the 300-1,800 m/z range at 70,000-mass resolution. Data dependent acquisition of tandem MS/MS spectra were obtained for the 12 most intense ions using high-energy collision dissociation with 17,500-mass resolution and a dynamic exclusion of 30 seconds. Raw MS data were analyzed by Maxquant (v.1.6.14.0) using default parameters and a *P. aeruginosa* protein sequence collected from Uniprot (accessed August 2020). A 1% false discovery rate was used at both peptide and protein levels. For quantification, intensity-based absolute quantification (IBAQ) was used, and all subsequent analysis was performed using Perseus (v1.6.5.0). Proteins were filtered to remove potential contaminants, followed by log_2_ transformation and imputation of missing values with intensity of zero.

Analysis of Ub linkage sites was performed following an *in vitro* ubiquitination reaction composed of 2 µM PUL-1, 125 nM E1, 2 µM E2, 37.5 µM Ub, 10 mM MgCl_2_, and 5 mM ATP in 25 mM sodium phosphate (pH 7.4), 150 mM, NaCl, 0.5 mM DTT. The reaction was incubated at 37 °C for 120 min and terminated by the addition of reducing SDS sample buffer. The samples were run into 4-12% SDS-PAGE gels and Coomassie stained. Gel bands corresponding to PUL1-Ub and diUb were excised and processed as above for mass spectrometry analysis.

### Immunoblotting

Protein samples were resolved by 4-12% Tris-Glycine SDS-PAGE (BioRad) and transferred onto 0.22 µm nitrocellulose membranes. Membranes were blocked with Tris-buffered saline containing 0.1% (w/v) Tween-20 (TBS-T) with 5% (w/v) non-fat dried skimmed milk powder at room temperature for 1 hr. Membranes were subsequently probed with indicated antibodies in TBS-T containing 3% (w/v) bovine serum albumin overnight at 4 °C. Horseradish peroxidase (HRP)-conjugated secondary antibodies in TBS-T were then blotted for 1 hr at room temperature, prior to visualization with ECL. Ubiquitin was probed with 1:1000 anti-ubiquitin primary antibody (clone Ubi-1, Millipore MAB1510). Biotin was probed with 1:1000 anti-biotin primary antibody (Bethyl Laboratories, A150-109A).

### Ub-DHA profiling of PUL-1

3 µM recombinant PUL-1 was mixed with 500 nM E1, 5 µM E2, 50 µM Biotin-Ub-DHA, 10 mM MgCl_2_, and 5 mM ATP in 50 mM HEPES (pH 8.0), 150 mM NaCl, 1 mM DTT. The reactions were incubated at 37 °C for 80 min and supplemented with 1 mM ATP every 20 min. To remove isopeptide- and thioester-linked Ub-DHA complexes, USP21 (0.5 µM) was added to the resulting sample along with 5 mM DTT and incubated for an additional 30 min at 37 °C. Reactions were then terminated by the addition of reducing SDS sample buffer.

### Acyl-CoA dehydrogenase activity assay

Acyl-CoA dehydrogenase assays were carried out using the DCPIP method, which measures the reduction of DCPIP as an electron acceptor downstream of PMS as an intermediate electron carrier. Assays were carried out under the following conditions: 50 mM HEPES-KOH buffer (pH 8.0), 50 µM FAD, 100 µg/mL DCPIP, 100 µg/mL PMS, 50 µM octanoyl-CoA lithium salt, and 1 µM enzyme. Reactions were performed at room temperature in technical triplicate and initiated by the addition of octanoyl-CoA lithium salt. DCPIP reduction was measured by the decrease in absorbance at 600 nm in clear, flat-bottom, 96-well plates.

### PUL-1 thioester trapping assay

A master mix containing 125 nM E1, 2 µM UBE2L3, 37.5 µM Ub, 2 µM PUL-1, and 10 mM MgCl_2_ was prepared in 25 mM sodium phosphate (pH 7.0), 150 mM NaCl. The reaction was then initiated by the addition of 5 mM ATP and incubated for 7 min (WT) or 20 min (C4S and C4A) at 37 °C. The reactions were terminated by the addition of nonreducing SDS sample buffer. Resulting samples were either left untreated, reduced with 10 mM DTT, or treated with 0.010 N NaOH.

### Bioinformatic analysis

PUL-1 orthologues were identified by PSI-BLAST and Phyre2 searches^66,67^. The sequences were aligned using Jalview Software and TCoffee^68,69^. Protein structural models were obtained from the AlphaFold Protein Structure Database^51^, analyzed for homology with DALI^52^, and visualized with PyMol (www.pymol.org).

### C. elegans growth conditions

C. elegans hermaphrodites were maintained on *E. coli* OP50 at 20 °C unless otherwise indicated. Bristol N2 was used as the wild-type control obtained from the *Caenorhabditis* Genetics Center (University of Minnesota, Minneapolis, MN). The bacterial strain *E. coli* OP50 was grown in LB broth at 37 °C.

### Quantification of intestinal bacterial loads

Animals were synchronized by placing gravid adults on modified nematode growth media (NGM) agar plates (0.35% instead of 0.25% peptone) containing *E. coli* OP50 for 2 hours at 20 °C. The gravid adults were removed, leaving the eggs to hatch and develop at 20 °C. For quantification of colony forming units (CFU), bacterial lawns were prepared by spreading 50 µL of overnight culture on the complete surface of 6 cm-diameter modified NGM agar plates. The plates were incubated at 37 °C for 12-16 hours and then cooled to room temperature for at least 1 hour before seeding with young gravid adult hermaphroditic animals. Animals were exposed to the *P. aeruginosa* lawns for 24 hours at 25 °C, after which the animals were transferred to the center of fresh *E. coli* plates for 30 min to eliminate bacteria stuck to their body. Animals were then transferred to the center of a new *E. coli* plate for 30 min to further eliminate external bacteria. Animals were finally transferred to fresh *E. coli* plates a third time for 10 min. Afterward, ten animals/condition were transferred into 50 µL of PBS plus 0.01% Triton X-100 and ground. Ten-fold serial dilutions of the lysates were seeded onto LB plates containing 5 µg/mL doxycycline with or without 200 µg/mL of carbenicillin to select for *P. aeruginosa* and grown overnight at 37 °C. Single colonies were counted the next day and used to calculate CFU per animal. Three independent experiments were performed for each condition.

### Quantification of intestinal lumen bloating

Synchronized young adult *C. elegans* hermaphrodites were transferred to modified NGM plates containing *P. aeruginosa* lawns and incubated at 25 °C for 24 hours. After the indicated treatment, the animals were anesthetized using an M9 salt solution containing 50 mM sodium azide and mounted onto 2% agar pads. The animals were then visualized using a Leica M165 FC fluorescence stereomicroscope. The diameter of the intestinal lumen was measured using Image J software. At least 10 animals were used for each condition.

### C. elegans killing assays

Bacterial lawns were prepared on modified NGM plates as indicated above. Synchronized young adult *C. elegans* hermaphrodites were transferred *P. aeruginosa* lawns and incubated at 25 °C. Animals were scored at the indicated times for survival and transferred to fresh pathogen lawns each day until no progeny was produced. Animals were considered dead when they failed to respond to touch and no pharyngeal pumping was observed. Each experiment was performed in triplicate (n = 90 animals).

### Quantification and statistical analysis

Statistical analysis was performed with Prism 7 (Graph Pad). The Kaplan Meier method was used to calculate the survival fractions, and statistical significance between survival curves was determined using the log-rank test.

## ACKNOWLEDGEMENTS

We thank David Komander (WEHI) and Rachel Klevit (UW) for sharing expression plasmids, and Leigh Knodler (U. Vermont), Brett Finlay (UBC), and John Rohde (Dalhousie U.) for sharing bacterial strains. We thank members of our laboratories and the Seattle Ub Research Group for helpful discussions. This work was facilitated, in part, by the PMedIC joint research collaboration between OHSU and the Pacific Northwest National Laboratory (PNNL), which is a multi-program national laboratory operated by Battelle for the DOE under Contract DE-AC05-76RL01830. This work was supported by the Laboratory Directed Research and Development Program at PNNL (ESN), the IARPA FunGCAT program (the funders had no role in the design or interpretation of the experiments) (JNA), Oregon Health & Science University (JNP), the Medical Research Foundation of Oregon (JNP), the NIH National Institute of Allergy and Infectious Diseases (R01AI156900 to AA and R21AI176089 to NK), and the NIH National Institute of General Medical Sciences (R37GM070977 to AA and R35GM142486 to JNP).

## AUTHOR CONTRIBUTIONS

Conceptualization, JNP; Investigation, CGR, SK, AJO, MED, GDW, and JNP; Resources, AK, NVK, PPG, MPCM, JEM, JNA, and AA; Writing – original draft, CGR and JNP; Writing – review & editing, all authors; Supervision, JNP; Funding acquisition, ESN, JNA, NK, AA, and JNP.

---

**Supplementary Figure 1:**
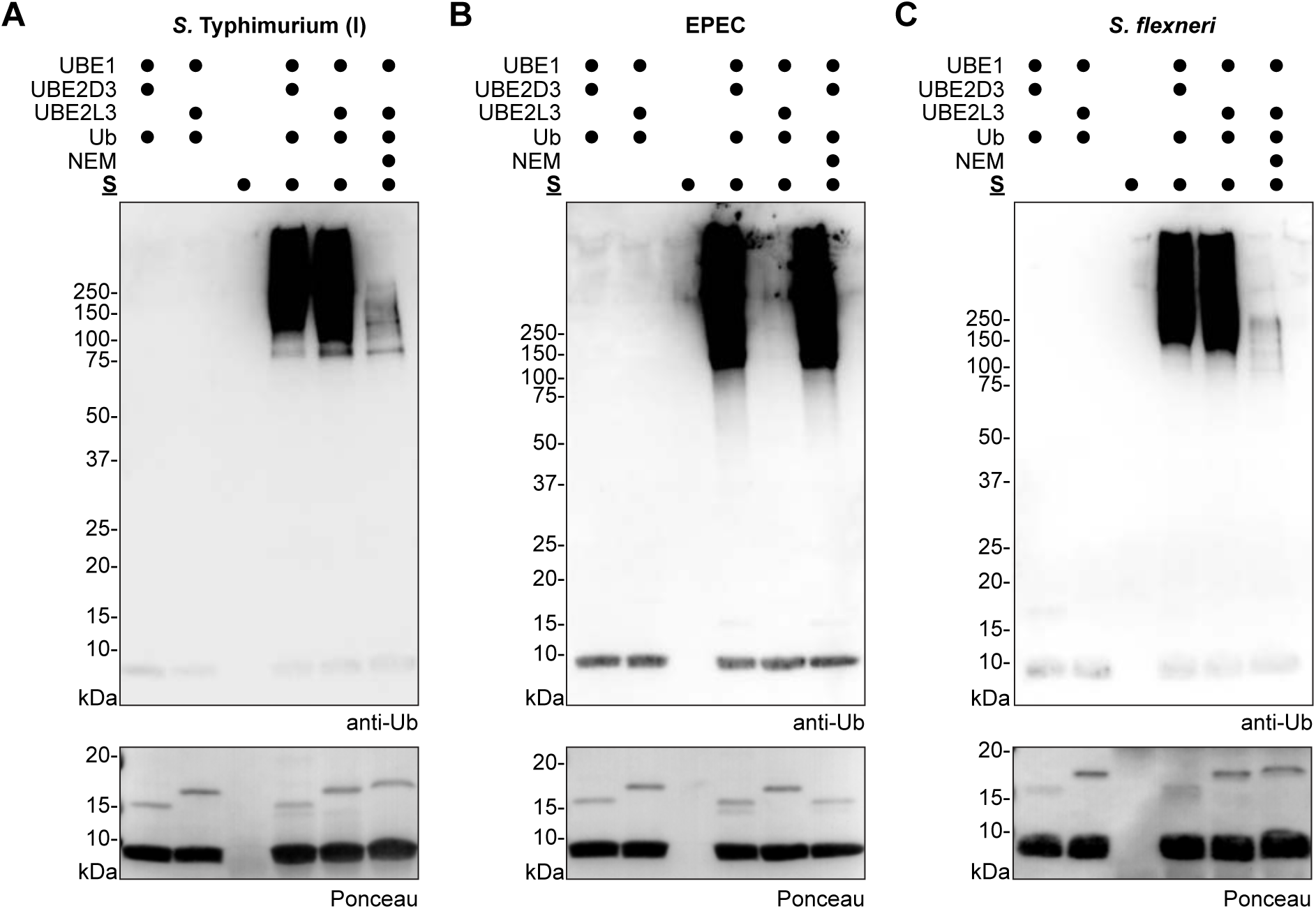
A functional screen for ubiquitin regulation. A. E3 ligase assays combining the indicated reaction components with the SPI-I secreted fraction from *S.* Typhimurium, with and without prior treatment with NEM. Reactions were resolved by SDS-PAGE and visualized by anti-Ub western blot. B. As in **A)**, for the secreted fraction from EPEC. C. As in **A)**, for the secreted fraction from *S. flexneri*.

**Supplementary Figure 2:**
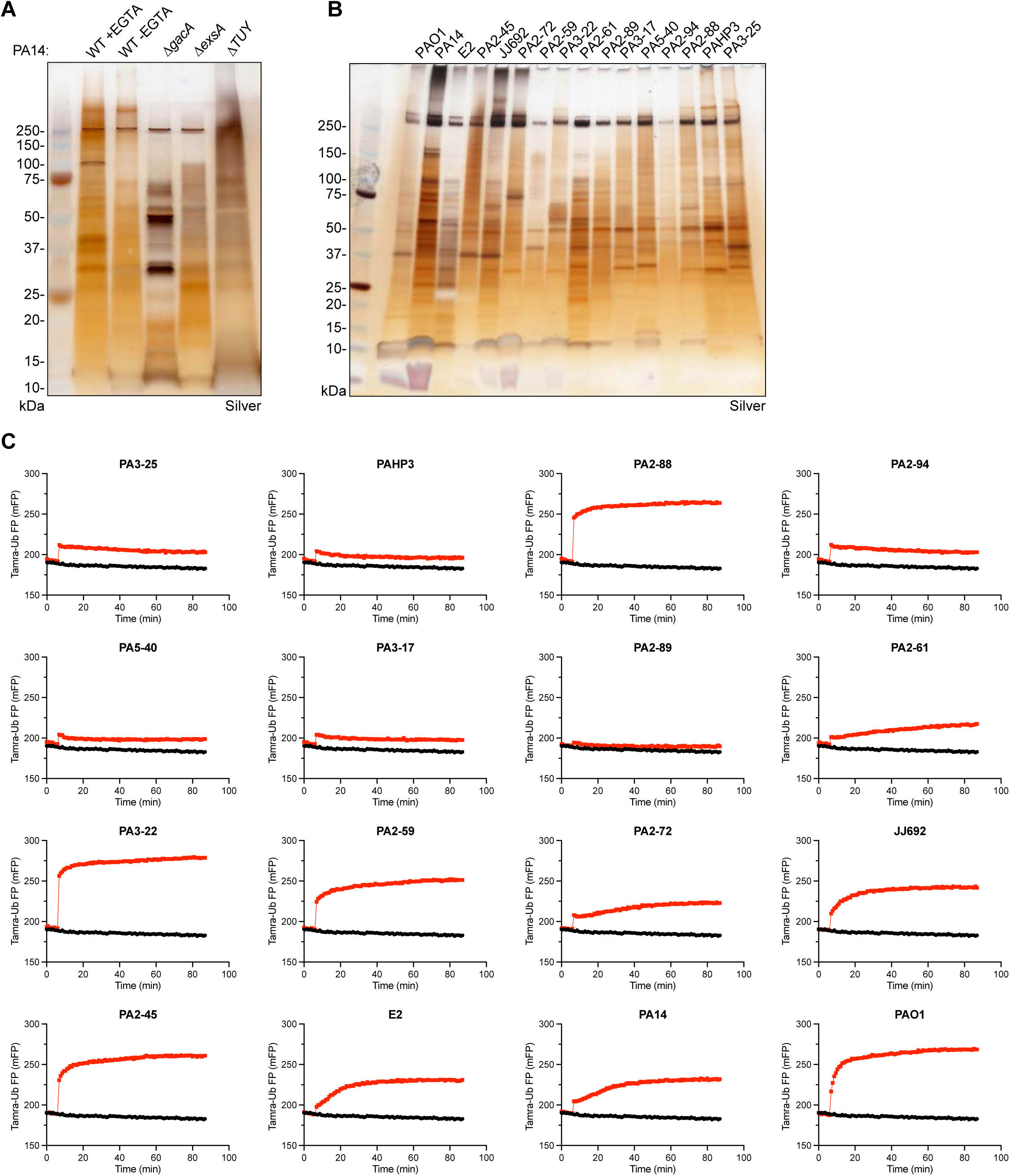
Detection of E3 ligase activity secreted by *P. aeruginosa*. A. Silver-stained SDS-PAGE analysis of secreted fractions generated from the indicated PA14 mutant strains, or in the absence of EGTA stimulation. B. Silver-stained SDS-PAGE analysis of secreted fractions generated from the indicated *P. aeruginosa* clinical isolates. C. Representative FP traces monitoring the Tamra-Ub ligase substrate following addition of secreted fractions from the indicated *P. aeruginosa* clinical isolates.

**Supplementary Figure 3:**
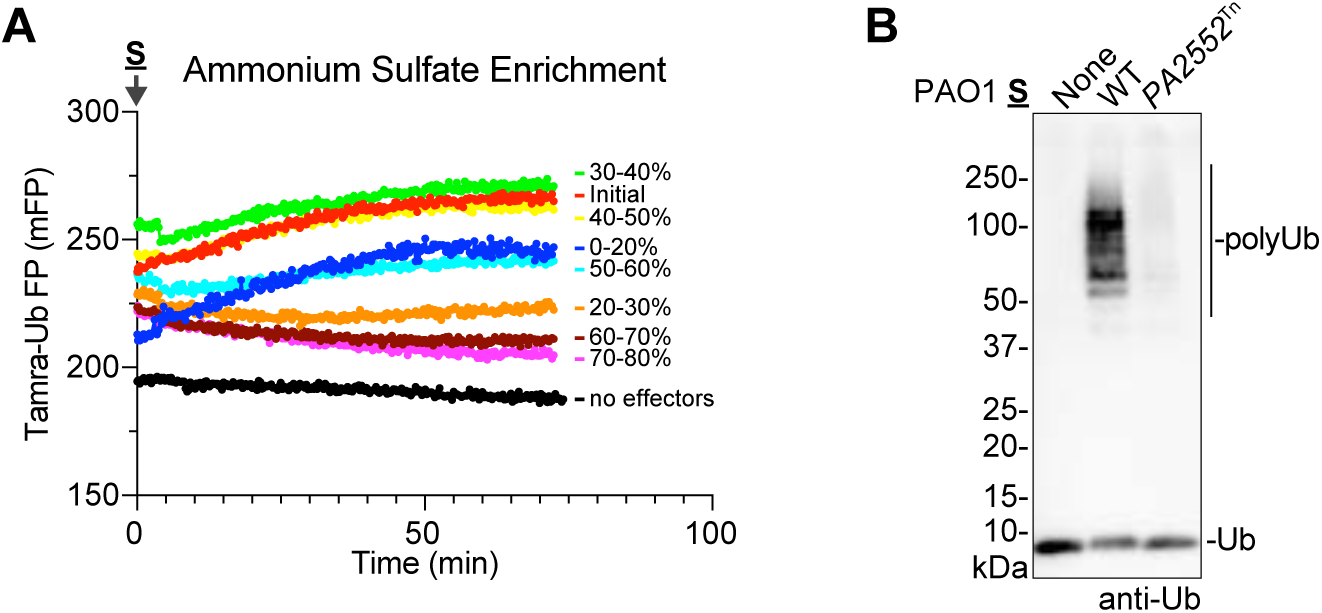
Identification of a *P. aeruginosa* E3 ligase. A. Representative FP traces monitoring the Tamra-Ub ligase substrate following addition of the indicated ammonium sulfate fractions of PA14 secreted protein. B. E3 ligase assays for secreted fractions generated from PAO1 wild-type or the *PA2552*^Tn^ mutant strain. Reactions were resolved by SDS-PAGE and visualized by anti-Ub western blot.

**Supplementary Figure 4:**
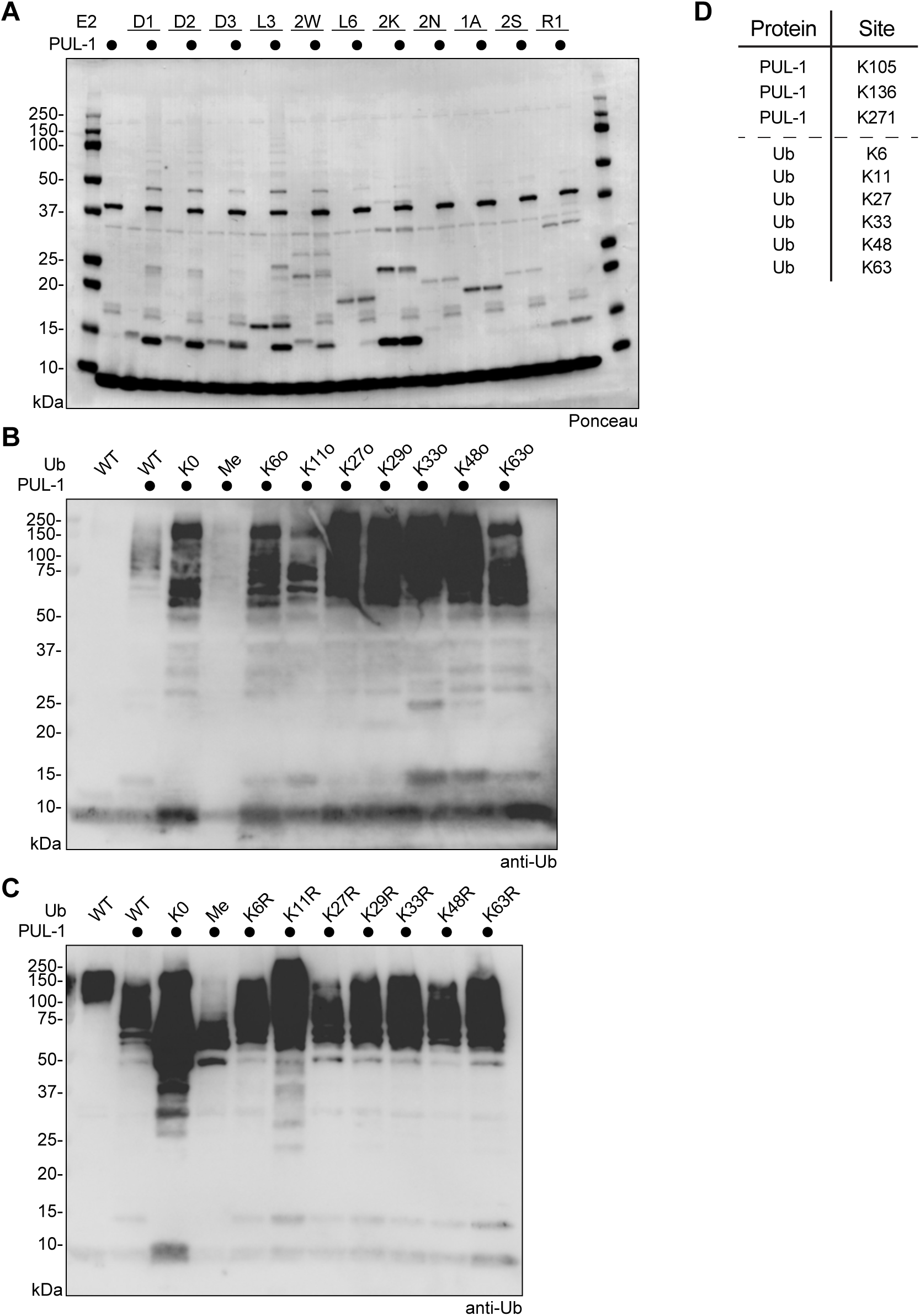
Characterization of PUL-1 E3 ligase activity. A. E3 ligase assays for recombinant PUL-1 and the indicated panel of E2 enzymes. Ponceau-stained visualization of Figure 4D. B. E3 ligase assays for recombinant PUL-1 and Lys-less (K0), methylated, or the indicated panel of K-only Ub mutants. Reactions were resolved by SDS-PAGE and visualized by anti-Ub western blot. C. E3 ligase assays for recombinant PUL-1 and Lys-less (K0), methylated, or the indicated panel of K-to-R Ub mutants. Reactions were resolved by SDS-PAGE and visualized by anti-Ub western blot. D. Ubiquitination sites identified by mass spectrometry following an *in vitro* PUL-1 ligase reaction.

**Supplementary Figure 5:**
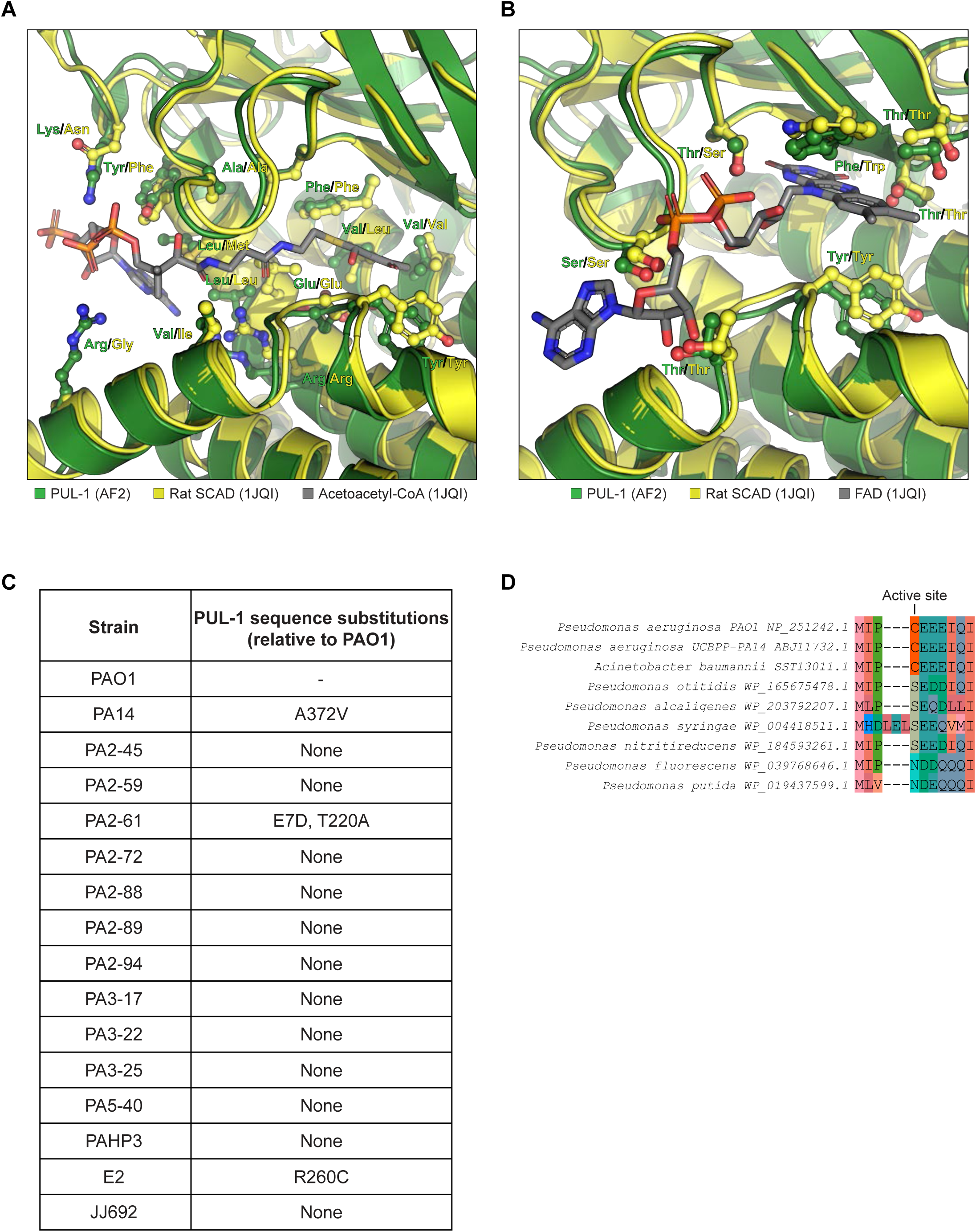
Structural analysis of the PUL-1 ligase fold. A. Structural overlay of the mitochondrial short-chain specific acyl-CoA dehydrogenase (SCAD) from rat (yellow, PDB: 1JQI), with acetoacetyl-CoA bound (grey sticks), and the PUL-1 AlphaFold2 model (green). Residues within the acyl-CoA-binding pocket are shown in ball-and-stick for both enzymes. B. Structural overlay of the mitochondrial short-chain specific acyl-CoA dehydrogenase (SCAD) from rat (yellow, PDB: 1JQI), with FAD bound (grey sticks), and the PUL-1 AlphaFold2 model (green). Residues within the FAD-binding pocket are shown in ball-and-stick for both enzymes. C. Conservation of PUL-1 orthologues among all *P. aeruginosa* clinical isolates presented in Figure 2F. Amino acid substitutions relative to PAO1 are listed. D. Sequence alignment of PUL-1 orthologues, focused on the region surrounding Cys4 of PAO1.

**Supplementary Figure 6:**
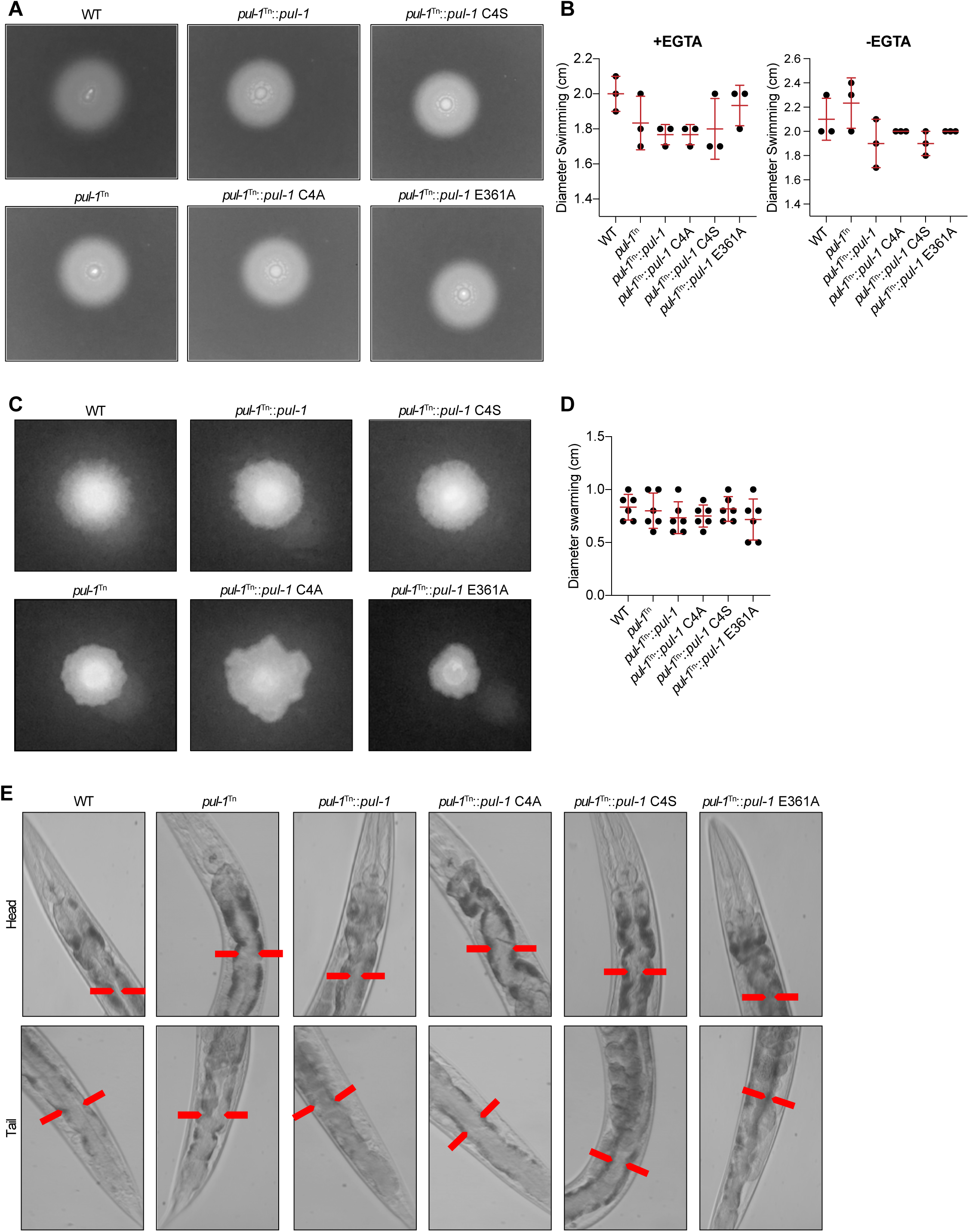
PUL-1 ligase activity modulates *P. aeruginosa* virulence. A. Representative images of *P. aeruginosa* swimming for WT PAO1 and the indicated *pul-1*^Tn^ mutant strains. B. Quantification of **A)**, under conditions with and without EGTA. Mean values and standard deviation are indicated in red. C. Representative images of *P. aeruginosa* swarming for WT PAO1 and the indicated *pul-1*^Tn^ mutant strains. D. Quantification of **C)**. Mean values and standard deviation are indicated in red. E. Representative images of *C. elegans* intestinal bloating near the head and tail, following infection with WT PAO1 or the indicated *pul-1*^Tn^ mutant strains. The intestinal lumen diameter is indicated by black arrows.

